# Membrane microdomains are crucial for *Mycobacterium marinum* EsxA-dependent membrane damage, escape to the cytosol and infection

**DOI:** 10.1101/2024.08.19.608731

**Authors:** Cristina Bosmani, Angélique Perret, Florence Leuba, Aurélie Guého, Nabil Hanna, Thierry Soldati

**Author notes:** These authors contributed equally (Cristina Bosmani, Angélique Perret).

## Abstract

Infection by pathogenic mycobacteria such as *Mycobacterium tuberculosis* disrupts the membrane of the Mycobacterium-Containing Vacuole (MCV). The key effector EsxA, secreted via the ESX-1 type-VII system, is pivotal in this process, yet its membranolytic activity is not fully elucidated. Infecting the amoeba *Dictyostelium discoideum* with *Mycobacterium marinum*, we demonstrate that the composition of the MCV membrane, notably its sterol-rich microdomains, significantly influences damage and rupture. Disruption of these microdomains through the knockout of organizing proteins, termed vacuolins, or through sterol depletion, markedly diminishes *M. marinum*-induced membrane damage and cytosolic escape, thereby increasing cellular resistance to infection. Furthermore, we establish that vacuolins and sterols are essential for the *in vitro* partitioning of EsxA within membranes. Extending our findings to mammalian cells, we show that the role of microdomain organizers and sterols is evolutionarily conserved; specifically, flotillin knockdown and sterol depletion enhance the resistance of murine microglial cells to *M. marinum* infection. Our results underscore the critical role of host membrane microdomains in facilitating mycobacterial membranolytic activity and subsequent cytosolic access, pivotal for a successful infection.

## INTRODUCTION

Tuberculosis, caused by *Mycobacterium tuberculosis* (Mtb), is a major global health issue, killing 1.3 million people in 2022^1^. Alveolar macrophages are the first line of defence against Mtb in the lungs, however Mtb manipulates the phagosome maturation pathway to establish a replicative niche inside a modified phagosome, the Mycobacterium-Containing Vacuole (MCV)^2,3^. *Mycobacterium marinum* (Mm), closely related to Mtb, causes a disease similar to tuberculosis in marine and freshwater vertebrates. Mm is widely used as a versatile experimental model for Mtb owing to its conserved virulence mechanisms, faster replication time and easier laboratory manipulation^4,5^.

*Dictyostelium discoideum* (Dd), a social amoeba, uses phagocytosis to feed on soil bacteria. Conservation of the phagosome maturation pathway^6^ and cell-autonomous defences with animal innate immune phagocytes, makes Dd a valuable model for studying host-pathogen interactions, particularly mycobacterial infections^7–12^. After uptake, innocuous bacteria are enclosed in a phagosome that matures to expose them to acidic pH, hydrolases, reactive oxygen species, and toxic metals necessary for bacterial killing^11,13–15^. In contrast, Mm and Mtb arrest phagosome maturation, creating an MCV from which they eventually escape in an ESX-1 (type VII secretion system)-dependent manner^12,16–18^.

Generation of MCV damage is crucial in the infection cycle. Mm-induced MCV damage starts as early as 15 minutes post-infection and largely depends on EsxA, a virulence factor secreted via ESX-1^8,19^. In both mammalian cells and Dd, the ESCRT and autophagy machineries are recruited to the damage site through ubiquitination of the damaged membrane, acting as a “repair-me” signal^12^. In Dd, TrafE, an E3 ubiquitin ligase, coordinates their recruitment, playing an important role in restricting the infection^20,21^. Damage accumulates later during infection, leading to bacterial escape to the cytosol and exposure to the xenophagy machinery^19–22^. Mtb and Mm lacking the ESX-1 secretion system and EsxA (ΔRD1 mutants) are unable to arrest phagosome maturation, induce less MCV damage, and are therefore strongly attenuated^16,16,22^. Scattered *in vitro* evidence suggests that MCV membrane composition is crucial for EsxA activity and might be assisted by lipidic factors such as phthiocerol dimycocerosates (PDIMs), which are complex branched cell-wall-associated lipids that reportedly induce sterol clustering in host membranes^23–25^.

In mammalian cells, membrane microdomains enriched in sterols and sphingolipids are stabilized by Flotillin-1 and -2. Flotillins are highly conserved microdomain scaffolding proteins at the plasma membrane and phagosome^26–28^. Microdomains are involved in infection by various pathogens, facilitating the activity of bacterial pore-forming toxins and cholesterol-dependent cytotoxins^29–31^. Flotillins also play roles in infections by pathogens like *Chlamydia pneumoniae*^32,33^ and *Anaplasma phagocytophilum*^34^.

Dd vacuolins (VacA, B, and C) are functional flotillin homologues, capable of oligomerizing and behaving as integral membrane proteins^35–37^. The absence of vacuolins impairs recognition and adhesion to particles, including Mm, affecting uptake but not killing of innocuous bacteria^37^. In this study we dissect the link between vacuolin microdomains and the biogenesis, maintenance, and repair of the MCV during Mm infection. We demonstrate that VacC is highly induced upon Mm-induced damage and that vacuolin-enriched membrane microdomains form at the MCV early post-infection. Disruption of these microdomains through vacuolin knockout and/or sterol depletion increases resistance to Mm intracellular growth and to LLOMe-induced sterile damage. Vacuolins and sterols are required for EsxA membrane partitioning *in vitro* and MCV damage *in vivo*. Finally, in murine microglial cells, sterol-rich flotillin microdomains also accumulate at the MCV and are important for successful infection.

## RESULTS

### Vacuolin C is Specifically Induced upon *M. marinum* Infection

To investigate whether mycobacteria manipulate vacuolin expression during infection, we analysed the transcriptomic response of wild-type (wt) Ax2(Ka) cells infected with GFP-expressing *M. marinum* (Mm) wt as described ^38^. The *vacC* gene, poorly expressed in vegetative cells (Dicty express^39^), was significantly induced as early as 1 hpi and remained high throughout the 48-hour infection cycle (Fig. 1a). *VacB* was transiently highly induced at 1 hpi, while *vacA* was upregulated at later time points (24-48 hpi).

**Figure 1:**
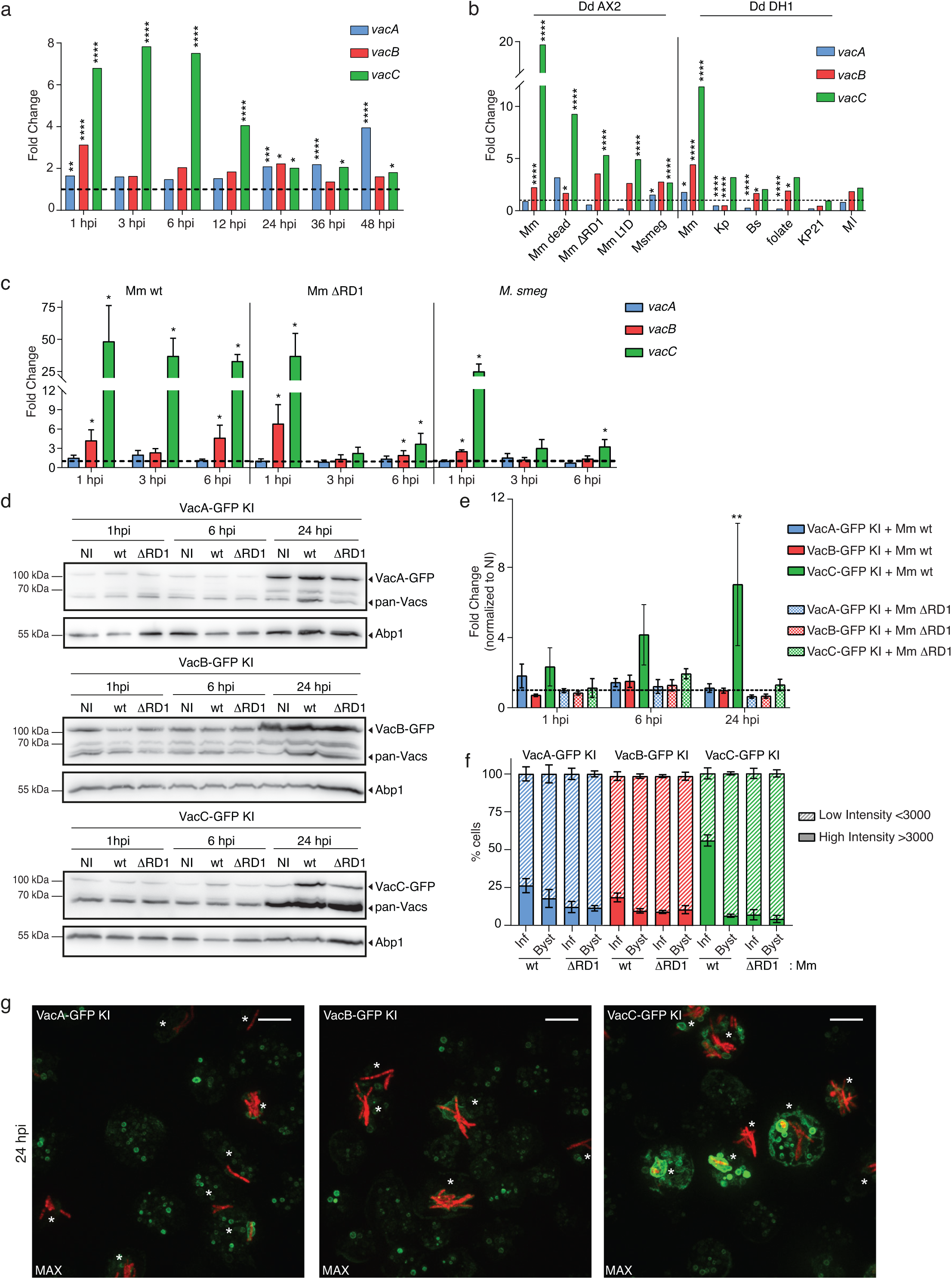
VacC is specifically induced upon *M. marinum* infection. **a.** RNA-sequencing of FACS-sorted wt cells infected with GFP-expressing Mm at different hours post infection (hpi). Fold change of each transcript compared to mock-infected cells (dashed line, N=3, *p≤0.05, **p≤0.01, ***p≤0.001, ****p≤0.0001). **b.** RNA-sequencing of AX2 or DH1 wt cells in contact with the indicated bacteria for 4 h, normalized and compared to mock-treated cells (dashed line) (N=3, *p≤0.05, ****p≤0.0001). Mm: *M. marinum*, Msmeg: *M. smegmatis*, Kp: *Klebsiella pneumoniae*, Bs: *Bacillus subtilis*, Ml: *Micrococcus luteus*. **c.** Quantitative RT-PCR of wt cell population infected with Mm wt or ΔRD1, or *M. smeg*. RNA levels were normalized to GAPDH and to mock-infected cells (dashed line, mean ± sem, N=4, *p≤0.05, Mann-Whitney test). **d.** Representative blots of lysates of a population of GFP knock-in (KI) cells infected with Mm wt or ΔRD1, or mock-infected (NI), immunoblotted with the indicated antibodies. **e.** Quantification of Vac-GFP bands normalized to Abp1 and NI cells (dashed line, N≥3, **p≤0.01, one-way ANOVA). **f.** Percentage of infected or bystander GFP KI cells with a low or high intensity GFP signal at 24 hpi (mean ± sem, N=2, n≥150 cells). **g.** Representative Max projections of indicated KI cell lines infected with Mm wt expressing mCherry at 24 hpi, the same settings were used to image all cell lines. *, infected cells; scale bar, 10 μm.

To determine whether *vacC* induction was specific to mycobacteria, we analysed the transcriptomic response of Ax2(Ka) and DH1 wt cells in contact for 4 hours with various Gram-positive and Gram-negative bacteria and mycobacteria strains (Fig. 1b,^40^). The *vacC* gene was significantly induced only when in contact with mycobacteria (Fig. 1b), correlating with the strain’s pathogenicity and capacity to induce MCV damage. Indeed, the highest induction was observed with Mm wt and the lowest with non-pathogenic *M. smegmatis*. RNAseq results were confirmed by qRT-PCR (Fig. 1c). The absolute fold change measured by qRT-PCR was higher than observed by RNAseq due to low basal expression of *vacC*. Unlike the constant upregulation of *vacC* with Mm wt infection, the ΔRD1 mutant and *M. smegmatis* only transiently induced *vacC* expression at 1 hpi.

To test whether VacC was correspondingly upregulated upon Mm infection, endogenous protein levels of each vacuolin were assessed by western blot using Vac-GFP knock-in strains (Vac-GFP KI^37^. VacC accumulated from 6 hpi and significantly at 24 hpi compared to mock-infected cells (NI; Fig. 1d-e), while no significant accumulation was observed for VacA or VacB. Additionally, infection with Mm wt, but not the ΔRD1 mutant, induced high VacC levels (Fig. 1d-e). Note that this analysis was performed on a population including infected and non-infected cells. High Content (HC) microscopy investigations at the single cell level confirmed higher VacC accumulation in Vac-GFP KI infected cells compared to non-infected bystanders (Fig. 1f-g). No significant difference in VacC-GFP levels was observed when cells were infected with the ΔRD1 mutant. These results demonstrate that VacC is a host reporter specific to Mm infection, likely triggered by EsxA-dependent MCV damage.

### Vacuolins Gradually Accumulate at the MCV Throughout Infection

Using Vac-GFP KI cells infected with mCherry-expressing Mm, we monitored vacuolin recruitment by live microscopy (Fig. 2a-b). Each vacuolin was present at the MCV as early as 1 hpi, with about 60% of MCVs coated with a vacuolin on the first day of infection (Fig. 2b). The association of vacuolins with the MCV changed over time, starting with a patchy distribution (Fig. 2a, c, e top panel) and progressing to coat the entire MCV, with about 75% of vacuolin-positive MCVs having a continuous vacuolin coat at 24 hpi (Fig. 2c, e middle panel). These results were confirmed using recombinant antibodies against endogenous VacA and VacB (Supplementary Fig. 1a). At late stages of infection, MCV damage became apparent, leaving only about 30-40% intact MCVs (Fig. 2d). The 3D reconstruction of deconvoluted live images of VacC-GFP KI infected cells highlighted the steps of vacuolin coat formation around the MCV (Fig. 2e). At 4 hpi, VacC-GFP forms patches around the MCV that extend to fully cover the MCV membrane at 24 hpi (Fig. 2e top and middle panels). At 24 hpi, Mm perforates the MCV and vacuolin coat to access the cytosol (Fig. 2e bottom panel). To monitor sterols at the MCV, we used filipin and the sterol sensor D4H*^41^ (Fig. 2f-g). While GFP-D4H* labelled the cytosolic leaflet of the MCV (Fig. 2f), filipin stained both the membrane and the MCV lumen (Fig. 2g). Moreover, VacC-GFP and mScarlet-D4H* co-localized at the MCV, providing strong evidence for sterol-rich microdomains (Supplementary Fig. 1b).

**Figure 2:**
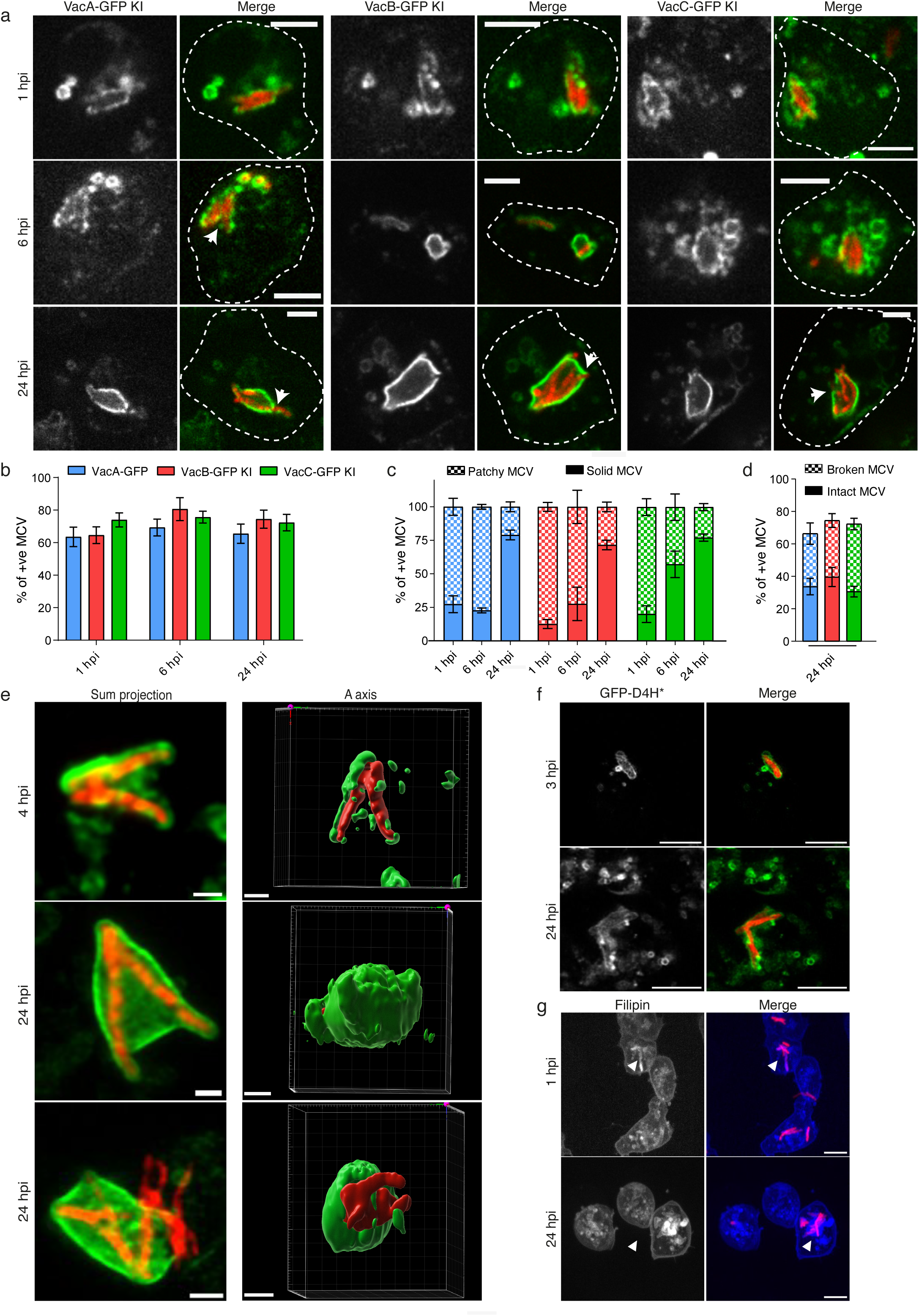
Vacuolin microdomains gradually accumulate at the MCV during infection. **a.** Representative images of indicated Vac-GFP KI cell lines infected with Mm wt expressing mCherry. Arrows, broken MCVs; scale bar, 5 μm. **b.** Quantification of a. presenting the proportion of MCVs positive for each vacuolin (mean ± sem, N=4, n≥200 MCVs). **c.** Quantification of a. presenting the percentage of MCVs showing a patchy or solid vacuolin-coat (mean ± sem, N=4, n≥200 MCVs). **d.** At 24 hpi, MCVs were separated into visibly broken or intact MCVs. **e.** Representative live images of VacC-GFP KI cell infected with Mm wt expressing mCherry and 3D reconstruction, Scale bar, 1 μm. **f-g.** Representative images of wt cells infected with mCherry-expressing Mm wt fixed at indicated time points. (**f.)** cells expressing GFP-D4H* or (**g.)** stained with filipin. Scale bar, 5 μm.

### Vacuolin Microdomains Accumulate at the *M. marinum*-Containing Vacuole

In analogy to flotillins, we hypothesized that vacuolins might be part of detergent-resistant membrane domains. To test this, Vac-GFP KI (Supplementary Fig. 2a) or Vac-overexpressing (OE; Supplementary Fig. 2b) cells were lysed in cold Triton X-100, and Triton-soluble (TSF) and -insoluble fractions (TIF) were separated by centrifugation. Then, TIF was floated by centrifugation on a step sucrose gradient (TIFF). Under basal conditions, only a small fraction of each vacuolin isoform, either as a GFP-tagged or endogenous version detected by a pan-Vac antibody, was present in TIFs (Supplementary Fig. 2a-b). Overexpression increased partitioning in microdomains (Supplementary Fig. 2b). Infection with Mm wt resulted in a larger fraction of VacC-GFP partitioned into ordered domains, with a two-fold enrichment in TIF at 18 hpi (Fig. 3a-b), in contrast to LmpB, a constitutive resident of ordered domains^42^. Additionally, extraction of sterols with methyl-beta-cyclodextrin (MβCD) strongly decreased vacuolin accumulation during infection (Fig. 3c-g and Supplementary Fig. 2c-h). Quantitative imaging showed a two-fold increase in VacC-GFP signal in infected cells at 24 hpi, which was abolished by MβCD treatment (Fig. 3d-e). Analysis at the single MCV level revealed an 80% decrease of VacC-GFP accumulation at the MCV after 16 hours of MβCD treatment (Fig. 3f-g). A similar trend was observed for VacA-GFP (Supplementary Fig. 2c-e) and VacB-GFP (Supplementary Fig. 2f-h). Overall, these results demonstrate that the MCV gradually acquires microdomain features characterized by vacuolin and sterol accumulation throughout the infection cycle.

**Figure 3:**
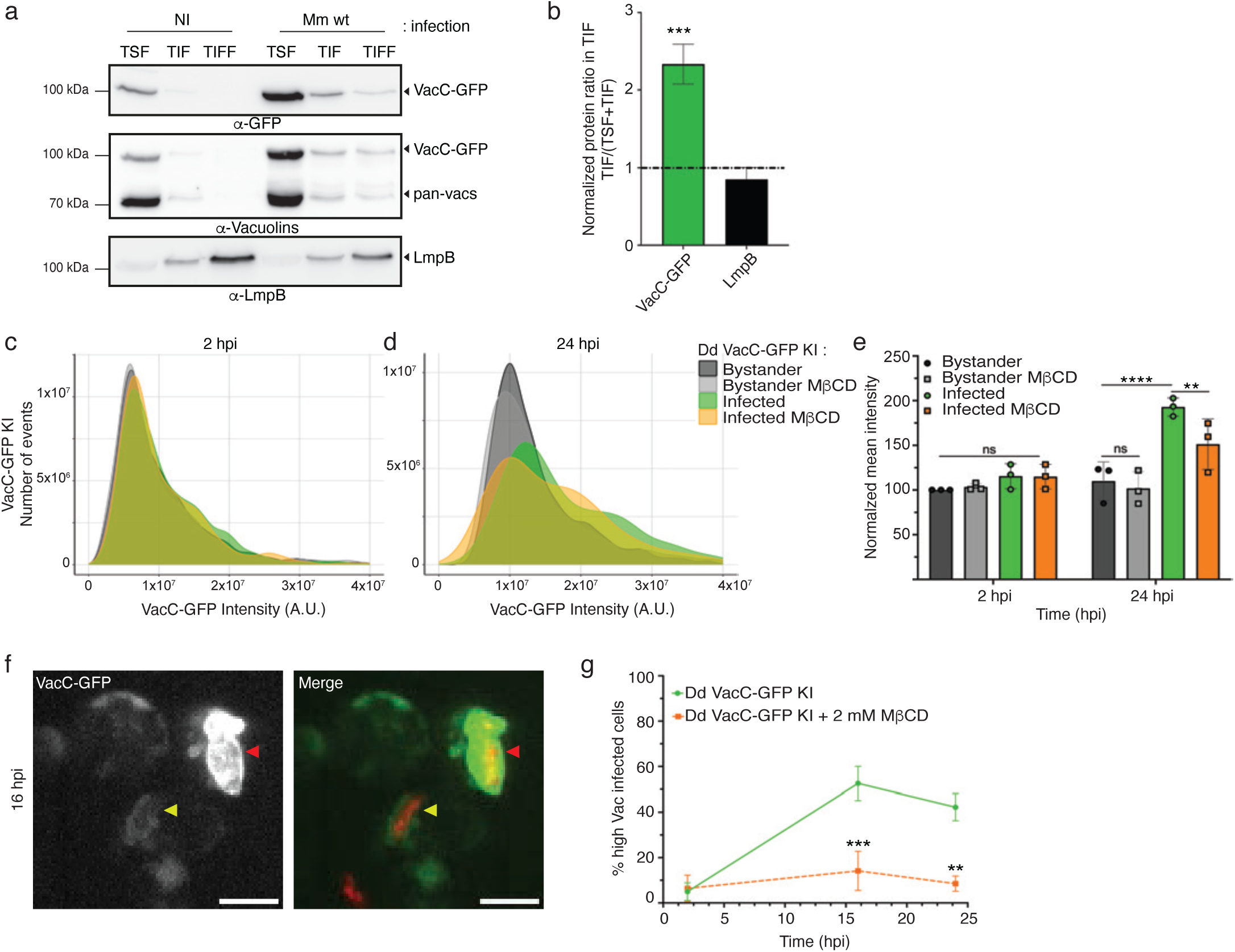
VacC microdomains accumulate at the MCV during infection. a-b. VacC-GFP KI cells were mock infected or infected with Mm wt and, after 16 hpi, lysed in cold Triton X-100. The Triton soluble (TSF) and insoluble (TIF) fractions were recovered, as well as the floating fraction (TIFF) from a sucrose gradient. Equal protein amounts of each fraction were loaded and immunoblotted with the indicated antibodies. a. Representative image of 3 independent experiments. b. Quantification of the fraction of VacC-GFP in the TIF compared to the total (TSF+TIF) (mean ± sd, N=3, one-way ANOVA ***p≤0.005). **c-g.** VacC-GFP KI cells infected with Mm wt expressing mCherry were monitored by HC microscopy. **c-d.** Normalized distribution of total VacC-GFP intensity per cell during infection and treatment with 2 mM of MβCD at 2 hpi (**c**) and 24 hpi (**d**) (N=3, n≥ 50000 cells). **e.** Mean intensity of **c.** and **d**. (mean + sd, N=3, two-way ANOVA, **p≤0.01, ****p≤0.0001). **f-g.** Analysis of **c-d.** at the MCV level. **f.** Representative image of MCV with high VacC-GFP intensity (red arrowhead) and low VacC-GFP intensity (yellow arrowhead). **g.** Quantification of high VacC-GFP MCVs (GFP intensity < 3000) after infection and treatment with MβCD (mean ± s.e.m, N=3, n≥150 cells, two-way ANOVA, **p≤0.01, ***p≤0.005).

### Disruption of Vacuolin Microdomains Confers Resistance to *M. marinum* Infection

To test whether the absence of vacuolins affects Mm virulence, we monitored the growth of wt and vacuolin mutant Dd strains on bacterial lawns containing mycobacteria. Whereas Dd wt does not grow well in the presence of Mm wt, cells with KOs of one or more vacuolins grew up to 10-fold better (Supplementary Fig. 3a-b), while growth in the presence of Mm ΔRD1 was unaffected. To directly examine the impact of vacuolin KO and sterol depletion, we measured intracellular Mm growth by monitoring either lux-expressing Mm in a plate reader (Fig. 4a-c) or GFP-expressing Mm by flow cytometry (Fig. 4d-e). In the absence of two (ΔvacBC) or all three vacuolins (ΔvacABC), Mm wt was not able to grow as efficiently as in wt cells, with a reduction of 50% (Fig. 4a and c) to 80% (Fig. 4d) at 72 hpi. Intracellular growth of the attenuated strain Mm ΔRD1, which induces only very limited MCV damage, is not significantly affected by vacuolin KO (Fig. 4a).

**Figure 4:**
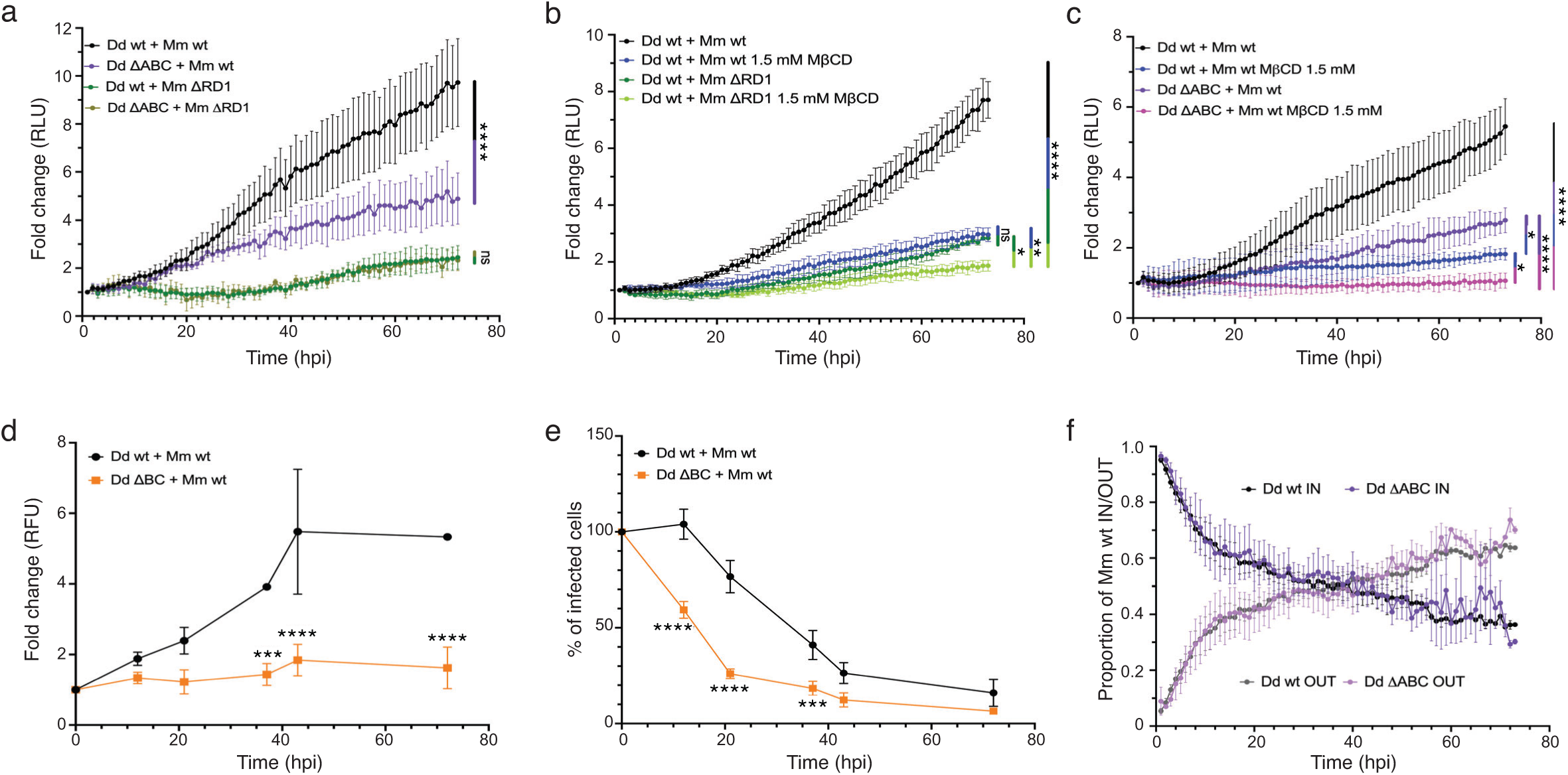
Disruption of vacuolin microdomains confers resistance to infection. **a-c.** Wt and ΔvacABC cells were infected with bioluminescent Mm wt or ΔRD1, and non-treated or treated with 1.5 mM of MβCD. Bioluminescence was measured for 72 hours in (mean fold change ± sem N=3, two-way ANOVA, *p≤0.05, **p≤0.01, ***p≤0.005, ****p≤0.0001). **d-e.** Wt and ΔvacBC cells were infected with GFP-expressing Mm up to 72 hours. At the indicated time points, fluorescence of infected cells (**d.**) and percentage of infected cells (**e.**) were measured by flow cytometry (mean ± sd, N=3, two-way ANOVA, *p≤0.05, **p≤0.01, ***p≤0.005, ****p≤0.0001). **g.** Wt and ΔvacABC cells were infected with GFP-expressing Mm wt and imaged by HC microscopy for 72 hours. After segmentation, the proportion of intracellular and extracellular bacteria calculated for each time point (mean fold change ± sem, N=3).

Sterol depletion with MβCD decreased Mm wt growth (Fig. 4b) in a dose-dependent manner (Supplementary Fig. 3c), reaching a reduction similar to the low growth level of the attenuated Mm ΔRD1 (Supplementary Fig. 3c). These doses of MβCD do not affect Mm growth in the medium (Supplementary Fig. 3d). Like the vacuolin KOs, MβCD only weakly impacted the intracellular growth of Mm ΔRD1 (Fig. 4b). Finally, sterol depletion in ΔvacABC cells produced a significant additive inhibition of intracellular growth of Mm wt (Fig. 4c). Our data indicate that vacuolins and sterols are crucial susceptibility factors, as their absence or depletion confers resistance to infection, and the combination abolishes Mm growth.

To gain further insight into the resistance mechanism, we investigated the dynamics of infection at the single-cell level using FACS and automated HC microscopy. In a standard infection, the proportion of infected cells decreases passively with time because they grow slower than uninfected bystander cells, and infection dissemination does not compensate. The doubling time of Mm is similar to that of the Dd host, both being around 8 hours^43,44^. Interestingly, the percentage of infected ΔvacBC cells decreased faster than wt cells, down to two-thirds compared to wt at 21 hpi (Fig. 4e), indicating active and early “curing” of the infection. This curing might be due to early pathogen release from the host cell, either by cell death, exocytosis, or non-lytic egress termed ejection^18,45^. To investigate whether interference with vacuolin microdomains is involved in early release, HC microscopy quantification of the proportion of extracellular versus intracellular Mm was performed in wt and ΔvacABC cells (Fig. 4f). Release of Mm wt from wt and ΔvacABC cells was observed with the same kinetics, with 50% of bacteria found extracellular at 27 hpi (Fig. 4f). These data suggest that early release is not a major mechanism to explain the resistance of vacuolin KO mutants to Mm wt infection.

### Vacuolin Microdomains are Important for LLOMe Sterile Damage Activity

Mm damages the MCV membrane in an EsxA-dependent manner, eventually resulting in escape to the cytosol^12,18,20^. Both increased retention in the MCV and precocious escape to the cytosol can result in reduced bacterial growth (Supplementary Fig. 4). The Mm ΔRD1 mutant is much weaker at both inducing damage and escaping from its compartment, causing a strong attenuation (Supplementary Fig. 4b1)^19^. Apparent attenuation can also be caused by host restriction. In cells with impaired MCV damage repair, Mm wt escapes precociously to the cytosol and is rapidly recaptured and restricted by xenophagy (Supplementary Fig. 4b2)^19–21^. We hypothesized that vacuolin microdomains modulate Mm escape from or retention inside the MCV by affecting the level of damage induced to the MCV.

We and others have shown that LLOMe (L-Leucyl-L-Leucine methyl ester), an endolysosome membrane disrupter, phenocopies the MCV damage induced by EsxA^21,46,47^. We used this tool to monitor the impact of microdomain disruption on endolysosome breakage. Using LysoSensor, a pH-dependent chemical probe accumulating in acidic compartments, we followed damage induced by LLOMe treatment (Supplementary Fig. 5). Wt cells lost LysoSensor signal faster and to a higher extent after LLOMe addition than ΔvacABC cells and took longer to recover the initial signal (30 min vs. 15 min), indicating a decrease in damage induced by LLOMe in ΔvacABC cells (Supplementary Fig. 5). Additionally, we monitored the recruitment of reporters of the ESCRT-mediated membrane-repair machinery (Fig. 5). Wt and vacuolin KO cells stably expressing Alix-GFP or GFP-Vps32 were pre-treated or not with 2 mM of MβCD and imaged by live HC microscopy. The single-cell analysis pipeline, including segmentation of cells and GFP-positive structures, is illustrated in Fig. 5c and h, resulting in the curves shown in Fig. 5d and i. Before the addition of LLOMe, all cells had a similar cytosolic signal (Fig. 5a and f), and instantaneously after LLOMe addition, puncta formed at sites of ESCRT-mediated repair, peaking at around 5-10 min for Alix-GFP (Fig. 5b and d) and at around 15-20 min for GFP-Vps32 (Fig. 5g and i), in good agreement with their successive roles in ESCRT recruitment and repair activity^48^. The LLOMe effect is transient, and cells fully repair and recover. Remarkably, both ΔvacBC and ΔvacABC cells, as well as cells treated with MβCD, showed less recruitment of ESCRT components and a faster disappearance of these structures, indicating a faster recovery (Fig. 5d and i). Quantification of the overall extent of reporter recruitment plotted in Fig. 5e and j indicated that the basal level of Alix-GFP and GFP-Vps32-positive structures is low and is maximally stimulated by LLOMe in wt cells (Fig. 5e and j). The reduction of both reporters in ΔvacBC and ΔvacABC cells is around 3 to 5-fold, values indistinguishable from treatment of wt cells with MβCD. MβCD treatment of vacuolin KO cells further significantly reduced the recruitment down to basal levels (Fig. 5e and j). In accordance with our LysoSensor observations, the reduction of ESCRT repair machinery indicates a strong reduction of LLOMe-induced damage, confirmed by intact cell morphology and the absence of cell death, contrary to the outcome of repair inhibition^21^. In conclusion, the absence of vacuolin and/or MβCD treatment renders the endolysosomal compartment more resistant to the lysosomotropic membrane disrupter LLOMe.

**Figure 5:**
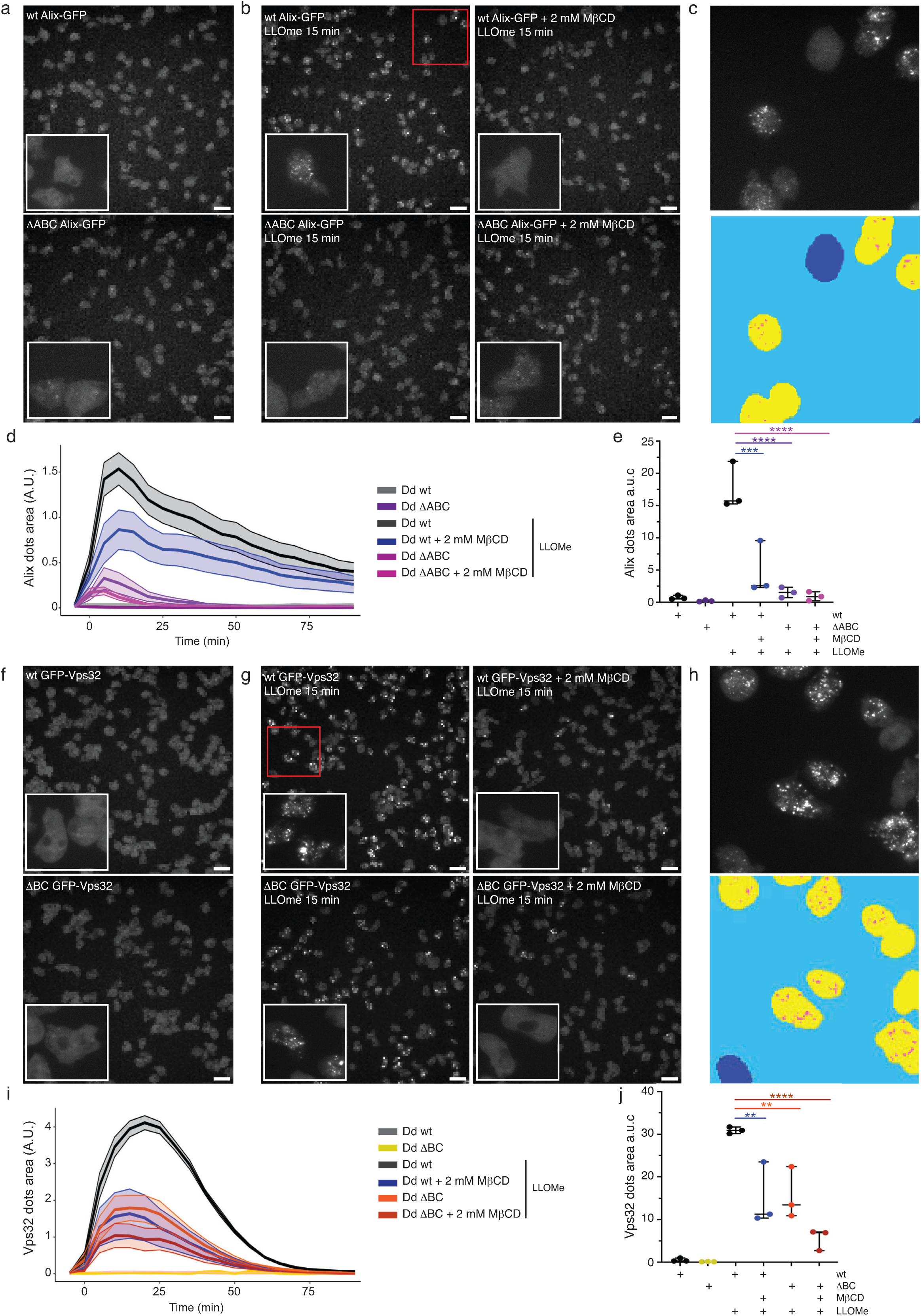
Disruption of vacuolin microdomains renders cells more resistant to LLOMe activity. **a-e.** Wt and 1′vacABC cells expressing Alix-GFP, treated or not with 2 mM of MβCD, were submitted to 4.5 mM of LLOMe. Representative live images of cells before treatment (**a.**) and 25 min after LLOMe addition, insets are magnified 3.5-fold (**b.**). Scale bar, 5 μm. **c**. Zoom of **b.** (red square) showing cell segmentation of non-reacting cells (blue), reacting cells (yellow) and Alix-GFP dots (pink) **d.** One representative experiment showing the area of Alix dots through time. The analysis is running per cell with <600 cells per condition. **e.** Area under the curve of **d.** (mean ± s.d, N=3, One-way ANOVA, ***p≤0.005, ****p≤0.0001). **f-j.** The same experiments and analysis were performed with wt and 1′vacBC cells expressing GFP-Vps32.

### Microdomain Disruption Impairs *M. marinum* Membranolytic Activity *In Vivo*

Given the above conclusion, we wondered whether sterol-rich microdomains are necessary for EsxA activity during infection. Ubiquitination of MCV and Mm surface proteins by the E3 ubiquitin ligase TrafE is one of the earliest hallmarks of damage at the MCV and is essential to trigger the recruitment of both the ESCRT and the autophagy machineries^20,21^. Once Mm reaches the cytosol, its surface becomes coated with the lipid droplet protein perilipin (mCherry-Plin;^49^). To monitor the steps of Mm-induced MCV damage and cytosol escape, we first quantified ubiquitin-labelling and GFP-Vps32 recruitment at MCVs in wt and vacuolin-KO cells, with and without MβCD treatment (Fig. 6a-b). Ubiquitin was found early around about 50% of Mm, whereas it decreased to around 35% at 5 hpi before levelled around 60% in wt and ΔvacBC. Treatment with MβCD significantly decreased it down to around 40% at 12 hpi and around 20% at 24 hpi (Fig. 6a). The early presence and decreasing association of GFP-Vps32 with the MCV in wt and ΔvacBC cells were similar to those of ubiquitin, and again were drastically affected in MβCD-treated cells, with 5% of infected cells showing Vps32 recruitment at 24 hpi compared to about 30% in non-treated cells (Fig. 6b). Compared to these reporters of damage and repair, translocation to the cytosol monitored by association with Plin is affected in all conditions compared to wt cells (Fig. 6c). At 1 hpi, only a minor fraction of Mm is exposed to the cytosol (around 15%). Starting from 5 hpi, Mm was increasingly associated with mCherry-Plin, reaching a plateau in wt and ΔvacABC cells around 45% and 35%, respectively (Fig. 6c). This apparent difference is just under the significance threshold, but the trend was observed in every replicate (Supplementary Fig. 6a). Strikingly, MβCD treatment decreases association with Plin in both cell types, with a plateau around 15-20% (Fig. 6c). Together, these results suggest that disturbing membrane microdomains by either knockout of vacuolins or sterol depletion limits damage to the MCV and severely inhibits cytosol translocation of Mm.

**Figure 6:**
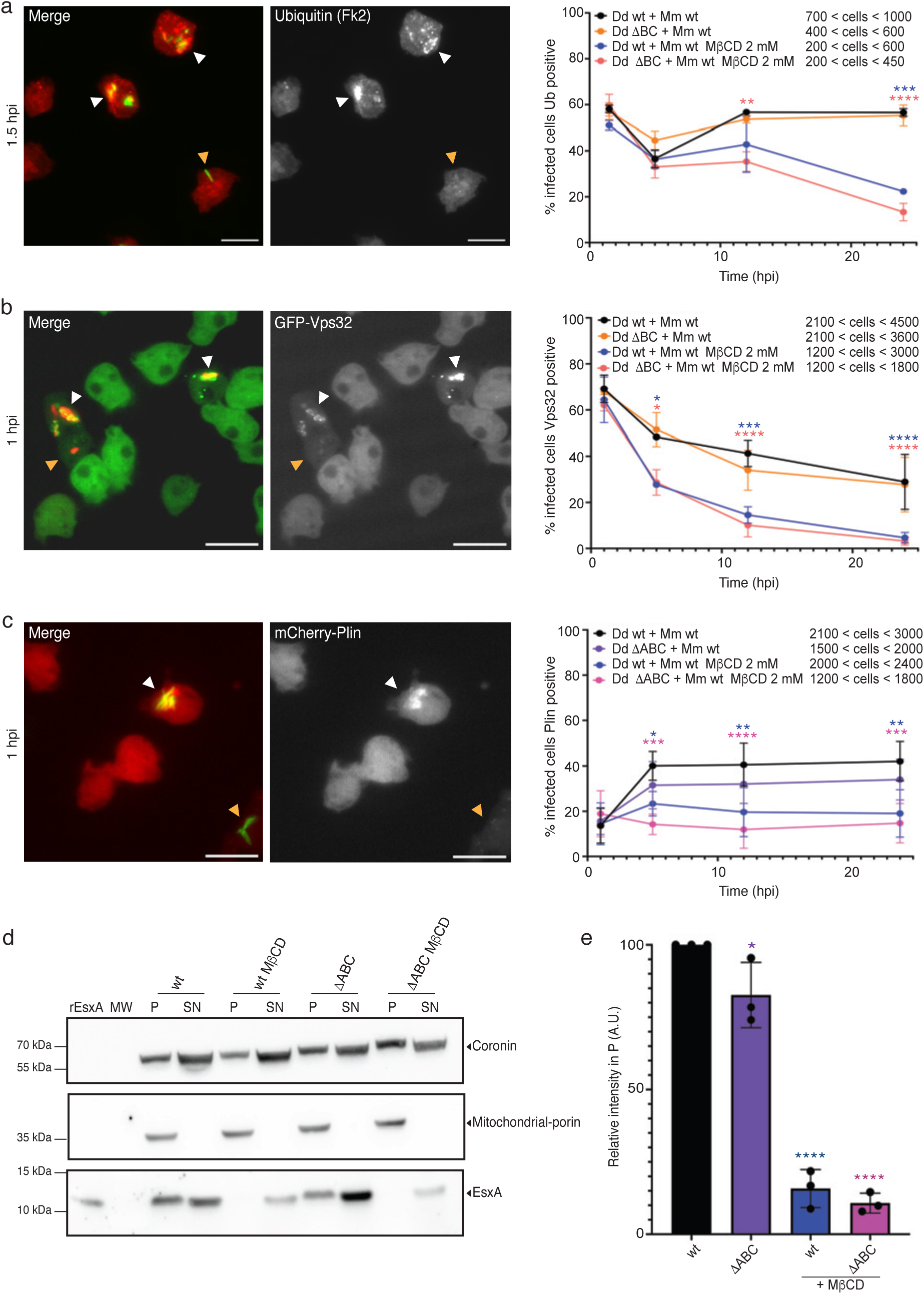
Microdomains are important for *M. marinum*-induced damage and ESX-A membranolytic activity. **a.** Wt and ΔvacBC cells were infected with BFP-expressing Mm wt, treated or not with 2 mM of MβCD, and, at the indicated time points, either immunostained for ubiquitin (FK2) or imaged live by HC microscopy. Left: representative image. Right: quantification of infected cells with ubiquitin labelling (mean ± s.d, N=3, two-way ANOVA, *p≤0.05, **p≤0.01, ***p≤0.005, ****p≤0.0001). **b-c.** The same experiment and analysis were performed with wt and 1′vacBC or 1′vacABC cells expressing GFP-Vps32 or mCherry-Plin respectively. **d-e.** Recombinant EsxA (rESAT-6) was incubated with post-nuclear supernatant (PNS) of wt or ΔvacABC cells, pre-treated or not with 10 mM of MβCD, followed by separation into supernatant (SN-cytosol) and pellet (P-membrane). Identical protein amounts were loaded and immunoblotted with the indicated antibodies. **d.** Representative western-bolt of 3 independent experiments. **e.** Quantification of the fraction of rEsxA in the membrane fraction normalized to mitochondrial-porin (mean ± sd, N=3, *p≤0.05, **p≤0.01, ****p≤0.0001).

### Microdomain Disruption Impairs EsxA Membrane Partitioning *In Vitro*

Mm induces membrane damage via ESX-1-mediated secretion of the small peptide EsxA, which inserts into and damages host membranes^17,50,51^, possibly with a preference for sterol-rich membranes^23,50,52^. To test this hypothesis, we monitored whether recombinant Mtb EsxA partitions into physiological host-derived membranes *in vitro* (Fig. 6d-e and Supplementary Fig. 6b-c). A whole membrane fraction was purified from wt Dd cells and incubated with rEsxA at different pH and with different membrane-to-EsxA ratios (Supplementary Fig. 6b-c). Optimal partitioning was observed at pH 6 (Supplementary Fig. 6b), corresponding to the pH of the MCV measured inside living Dd using Mm wt coated with FITC and TRITC (Supplementary Fig. 6d). As a control, the pH around an avirulent Mm L1D mutant^45^ that traffics to phagolysosomes is much lower (Supplementary Fig. 6d). We incubated rEsxA with membranes extracted from wt and ΔvacABC cells treated with or without 10 mM of MβCD. We observed a reduction of about 15% of rEsxA association with membranes purified from ΔvacABC mutants (Fig. 6d-e) and a drastic reduction of partitioning into membranes after sterol depletion, down 80% for wt membranes and 90% for membranes from ΔvacABC cells. Importantly, in accordance with the literature, we did not observe association of rEsxB with the membrane fraction under any condition (Supplementary Fig. 6e). Our results indicate that microdomains enriched in sterols and vacuolins are important for EsxA partitioning into host membranes, likely potentiating its membrane-damaging activity.

### Flotillin Microdomains are Important for *M. marinum* Infection in Murine Microglial BV-2 Cells

To directly probe whether microdomain involvement is conserved in animal phagocytes, we monitored BV-2 cells stably expressing Flot2-GFP or GFP-D4H* infected with mCherry-expressing wt Mm (Fig. 7a). Flot2 and the genetically encoded sterol probe D4H* localized around the bacteria, suggesting, like in Dd, an enrichment of sterol-rich microdomains at the MCV. Additionally, filipin staining of infected GFP-D4H* cells confirmed the presence of sterol at the MCV membrane and in its lumen (Fig. 7b).

**Figure 7:**
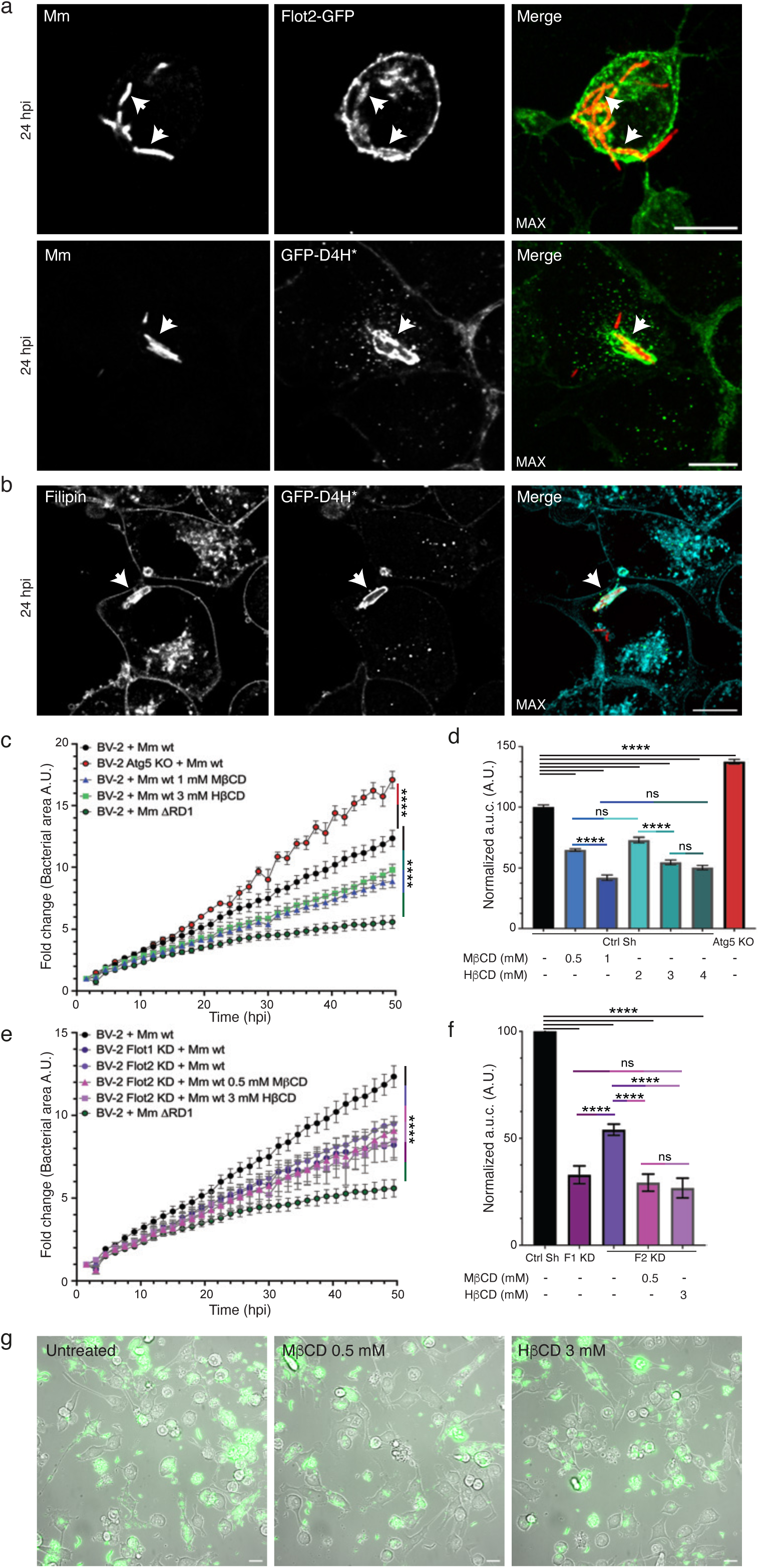
Disruption of flotillin microdomains confers resistance to infection in murine BV-2 microglial cells. **a-b.** BV-2 cells expressing Flot2-GFP or GFP-D4H* were infected with mCherry-expressing Mm wt, fixed at 24 hpi and stained with filipin. Scale bar, 10 μm. **c-g**. BV-2 cells were infected with GFP-expressing Mm wt and monitored by HC microscopy for 48 hours. **c. and e**. Growth curves of Mm measured as the area sum of the GFP signal at every time point (mean fold change ± sem N=3, two-way ANOVA, *p≤0.05, **p≤0.01, ***p≤0.005, ****p≤0.0001). **d. and f**. Analysis of the area under the curve of **c.** and **e.** normalized to Mm wt (100 %) and ΔRD1 (0 %) (mean ± sd N=3, one-way ANOVA, *p≤0.05, **p≤0.01, ***p≤0.005, ****p≤0.0001). **g.** Representative live images of **c**. at 33 hpi. of non-treated and cyclodextrin conditions. Scale bar, 5 μm

To better dissect the role of microdomains, we quantified intracellular Mm growth by HC microscopy in BV-2 cells. We first validated this BV-2 infection model by comparing Mm wt growth in wt cells and Δatg5. As expected, Mm wt grows better in host cells in the absence of autophagy restriction (Fig. 7c and e)^53,54^. Similar to the Dd infection model, the intracellular growth of Mm ΔRD1 is strongly attenuated (Fig. 7c and e). Treatment with MβCD or 2-hydroxypropyl-β-cyclodextrin (HβCD) also resulted in a dose-dependent decrease of intracellular Mm growth (Fig. 7c-d). We generated stable BV-2 cell lines with knock-down (KD) of Flot-1 and -2, the vacuolin orthologues, and monitored Mm infection (Fig. 7e-f and Supplementary Fig. 7a). Compared to the control (Ctrl Sh), KD of Flot-1 and -2 reduced Mm intracellular growth (Fig. 7e-f), similar to the decrease observed with CD treatments. We also observed an additive effect of CD treatment during infection of Flot-2 KD cells (Fig. 7e-f). Interestingly, CD treatments appeared to constrain the spreading of Mm wt inside BV-2 cells (Fig. 7g and Supplementary Fig. 7b), as revealed by measuring the area of all microcolonies (Supplementary Fig. 7c-h). The microcolony area increased under CD treatment (Supplementary Fig. 7e and g), indicative of decreased membrane damage resulting in confinement in the MCV, similar to that of the ΔRD1 Mm mutant (Supplementary Fig. 7e-g). CD treatment increased microcolony area in a dose-dependent manner and to a similar extent as Flot-1 and -2 KD. They also have an additive effect (Supplementary Fig. 7e and h). Our results in microglial cells demonstrate an important role of sterol-rich flotillin microdomains during Mm infection, highlighting the extreme evolutionary conservation of basic mechanisms of mycobacterial virulence and host cell-autonomous immunity.

## DISCUSSION

In *D. discoideum* (Dd), vacuolins are integral membrane proteins that oligomerize and define specific microdomains of the plasma and phagosomal membranes, similar to their mammalian homologues, flotillins^37^. Here, we propose that in evolutionarily distant phagocytes, host membrane microdomains are hijacked by pathogenic *M. marinum* (Mm) and are required for the efficient membranolytic activity of EsxA, resulting in membrane damage and escape from the MCV (Fig. 8).

**Figure 8:**
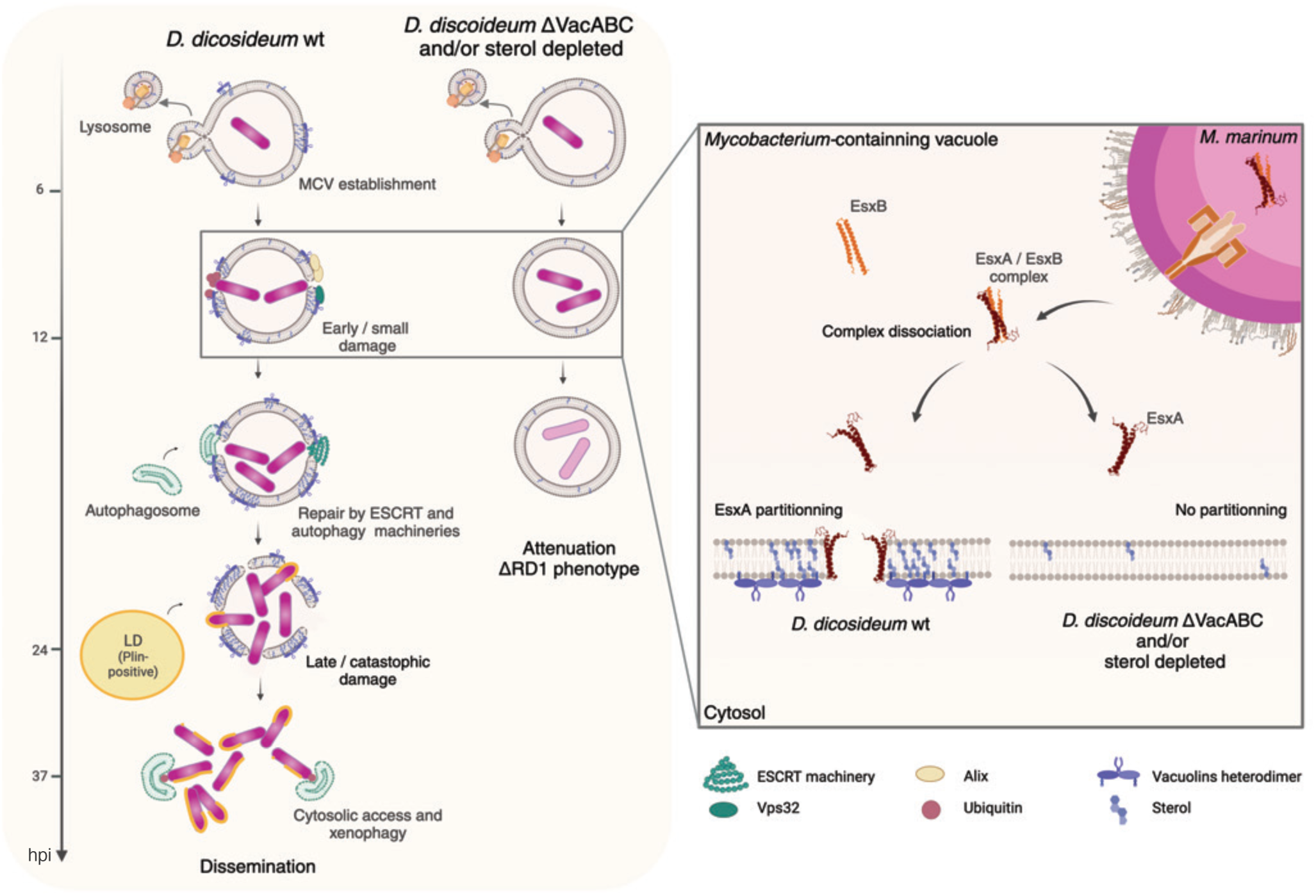
Working model of the role of membrane microdomains during Mm infection. Briefly, membrane microdomains accumulate at the MCV during infection, facilitating EsxA partitioning into the MCV membrane, and subsequent damage and Mm escape to the cytosol. Disruption of microdomains inhibits partitioning of EsxA into the MCV membrane, thereby limiting damage and leading to an attenuated infection.

Vacuolins, particularly VacC, are highly induced both at the mRNA and protein levels during infection with Mm wt, but to a lower extent with attenuated mycobacteria such as the ΔRD1 mutant and *M. smegmatis* (Fig. 1). The early induction of VacC is likely triggered by common mycobacterial pathogen-associated molecular patterns (PAMPs). However, sustained VacC expression requires mycobacteria with functional ESX-1, suggesting a specific response to damage caused by the secretion of membranolytic factors (Fig. 1c-f).

Vacuolins are initially present at the MCV in patches, and as the infection progresses, the bacterium becomes surrounded by a densely vacuolin-coated MCV, allowing clear visualization of catastrophic MCV damage (Fig. 2). Vacuolins are functional homologs of flotillins, and like them, are involved in organizing sterol-rich ordered domains. Indeed, vacuolins are present in Triton-insoluble fractions at steady state in uninfected cells (Supplementary Fig. 2a-b), and Mm infection significantly increases this partitioning (Fig. 3a). This biophysical evidence corroborates the transition from a patchy to a solid MCV coat (Fig. 2). Additionally, sterols also concentrate at the MCV during infection (Fig. 2f-g). Most importantly, sterol depletion with MβCD leads to a drastic decrease in vacuolin accumulation at the MCV (Fig. 3c-g). Altogether, these data indicate a specific accumulation of both microdomain components at the MCV, a phenomenon amplified for Mm with MCV-damaging activity. This emphasizes that over time, the lipid and protein composition of the MCV undergoes significant changes. MCV proteomic analyses confirm that virulent Mm profoundly impact MCV composition compared to non-pathogenic or attenuated mycobacteria, with, for instance, an important decreased abundance of proteins related to killing intracellular bacterial and a higher abundance of proteins linked to lipid acquisition, as well as ESCRT and autophagy machineries^55^.

We have previously described a role for VacB as a susceptibility factor involved in Mm infection^8^. However, because the Dd mutant used in that study was an accidental vacuolin B and C double KO^37^, here we present a more complete study and dissection of the roles of all three vacuolins and of sterols in the establishment of the Mm replicative niche. We confirm that the absence of VacB, as well as VacA and/or VacC, confers resistance to infection with Mm (Fig. 4). This phenotype was also observed to a similar extent in cells depleted of sterols using MβCD. Importantly, we found that growth of the ΔRD1 Mm mutant is similarly affected in wt cells, vacuolin KO mutants and sterol-depleted cells (Fig. 4a-b). In other words, the absence of vacuolins and sterol depletion specifically impact the growth of Mm wt but not of the ΔRD1 Mm mutant. This corroborates the hypothesis that vacuolin microdomains are specifically hijacked by Mm and involved in the establishment of the Mm niche in an ESX-1 and membrane damage-dependent manner. The interplay between sterols and vacuolins was emphasized by the observation of a stronger Mm wt growth defect in vacuolin knock-out cells treated with MβCD (Fig. 4c). We showed that the proportion of infected vacuolin KO cells decreases faster during the infection compared to wt cells, not because of earlier egress or release, but by an active restriction process (Fig. 4f). Altogether, these data indicate that disrupting membrane microdomains alters the escape of Mm from the MCV, resulting in an attenuated infection similar to that of the ΔRD1 mutant.

We wondered how vacuolin microdomains contribute to Mm escape from the MCV. Previous studies showed a decrease in Mtb uptake and growth inside macrophages after CD treatment, presumably by inducing cell apoptosis and bacterial death^56,57^. In our system, the addition of CD after uptake of Mm allowed us to specifically study the involvement of membrane microdomains during the intracellular life of Mm. Moreover, under our conditions, we observed neither increased cell death after treatment with MβCD nor earlier bacterial release, thus hypothesizing that less damage would be produced at the MCV in the absence of sterol-rich vacuolin microdomains.

We and others use LLOMe as a proxy to understand how mycobacteria-induced MCV damage is sensed and repaired by the host cell^21,46,47^. Using LysoSensor to follow endo-lysosomal leakage and two markers of damage sensing and repair, Alix and the ESCRT-III subunit Vps32^20,21^, we showed that the presence of sterols and microdomain organizers is required for LLOMe to induce endo-lysosomal damage. This represents the first direct evidence that membrane composition affects the action of this lysosomotropic membrane disrupter (Fig. 5 and Supplementary Fig. 5). Vacuolin microdomains also play a role in facilitating Mm-induced MCV damage. We showed that the levels of ubiquitination in the vicinity of the bacterium and Vps32 recruitment, which are among the earliest signs of damage and repair, are lower under MβCD treatment, suggesting that sterols are important for the induction of early and/or small damage (Fig. 6a-b). In addition, Plin binding to Mm, a hallmark of extensive catastrophic damage and cytosolic access, is strongly affected in sterol-depleted cells but also appears to be reduced in vacuolin KO cells (Fig. 6c and Supplementary Fig. 6a). This evidence indicates two distinct stages in MCV damage progression, and that interference with cytosol escape leads to a confinement that is sufficient to strongly attenuate Mm intracellular growth (Fig. 4a).

Mm and Mtb secrete EsxA, which has long been described as a membrane-disrupting toxin^17,50,58,59^. EsxA is secreted together with EsxB, its putative chaperone, through the ESX-1 secretion system. To damage the membrane, EsxA first needs to dissociate from EsxB, which occurs at low pH and was recently shown to require acetylation of EsxA^50,60^. During Mm infection of Dd, the average MCV pH remains around 6 (Supplementary Fig. 6d), probably due to early membrane damage leading to proton leakage. Additionally, we previously documented that some Mm experience a transient drop in pH, resulting in a fraction of MCVs being LysoSensor-positive at an early stage of infection^19^. We assume that a slightly acidic pH allows the dissociation of EsxA from its chaperone. Moreover, partitioning of EsxA in the MCV membrane strongly depends on the presence of vacuolins and the level of sterols (Fig. 6d). We propose that the progression of damage observed at the MCV mirrors the accumulation of vacuolins and sterol-rich microdomains. Interference with microdomains phenocopies the attenuation of the ΔRD1 mutant. Other additional virulence factors, such as PDIMs, reported to induce sterol clustering^23–25,61^, likely also participate in creating the appropriate membrane environment for EsxA activity.

We showed that the involvement of membrane microdomains during Mm infection is not restricted to Dd but is also observed in mammalian phagocytes. Flotillin homologues of vacuolins and sterol reporters are also recruited to the MCV in BV-2 microglial cells (Fig. 7a-b). Interestingly, similarly to Dd, D4H* and filipin co-localize at the MCV, but filipin staining was also enriched inside the MCV, pointing to a sterol accumulation not only at the MCV membrane but also inside, which might serve as a carbon source for Mm^49,62,63^. CD treatment and flotillin KD also render BV-2 cells more resistant to Mm infection (Fig. 7), apparently by constraining Mm dissemination in the cells. Again, as in Dd, CD treatment of Flot2-KD cells amplifies resistance to the infection (Supplementary Fig. 7).

In this study, we propose that Mm controls the composition and properties of the MCV membrane by manipulating host membrane microdomains proteins and lipids, thus enhancing EsxA partitioning and membranolytic activity, allowing MCV escape and Mm survival inside distantly evolutionarily related phagocytes, such as the amoeba D. discoideum and mammalian microglial cells (Fig. 8).

## ACKNOWLEDGMENTS

We gratefully acknowledge Dr. P. Cosson (University of Geneva) and Dr. J. King (University of Sheffield) for their invaluable discussions and suggestions, and Dr. P. Brodin (Institut Pasteur de Lille) for her insights on using cyclodextrin in mammalian cells. We also appreciate the support from the staff at the Photonic Bioimaging Center, the FACS Core Facility, and the iGE3 Genomics Platform at the Faculty of Sciences and Faculty of Medicine, University of Geneva. Special thanks to Dr. D. Moreau and the ACCESS Geneva Imaging Facility for their assistance with high-content microscopy experiments. We acknowledge the receipt of reagents from BEI Resources, NIAID, NIH, including ESAT-6 and CFP-10 recombinant protein standards, and polyclonal anti-*Mycobacterium tuberculosis* CFP10 and ESAT6. This work was supported by Swiss National Science Foundation grants 310030_188813 and 310030_219364. TS is a member of iGE3 (http://www.ige3.unige.chhttp://www.ige3.unige.ch).

## AUTHOR CONTRIBUTIONS

Every author made substantial contributions to the study, are accountable for the accuracy and integrity of the work and approved the submitted version of the manuscript. CB and AP contributed equally to all aspects of the work and manuscript, conception of the project and experimental design, the acquisition, analysis, and interpretation of data, and drafted the work and implemented revisions. FL, AG and NH participated in preliminary and/or contributing experimental work and performed the accompanying analysis and representation. TS acquired the funding, participated in the conception of the project and the design of the experimental strategies, the analysis and representation of the results, revised and edited the manuscript.

## MATERIALS AND METHODS

### *D. discoideum* Strains Culture and Plasmids

- **Culture Conditions:** *D. discoideum* strains and plasmids are listed in Extended Table 1. Cells were axenically grown at 22°C in HL5c medium (Formedium) supplemented with 100 U/mL of penicillin and 100 μg/mL of streptomycin (Invitrogen). Plasmids were transfected into Dd by electroporation and selected with the relevant antibiotic. Hygromycin was used at a concentration of 15 μg/mL for KO cell lines and 50 μg/mL for cell lines with reporters integrated at the safe-haven act5 locus. Blasticidin was used at a concentration of 5 μg/mL for KI and KO cell lines.

### BV-2 Strains Culture and Plasmids

- **Culture Conditions:** BV-2 strains and plasmids are listed in Extended Table 1. Cells were grown at 37°C in DMEM complemented with 10% FBS, 100 U/mL of penicillin, and 100 μg/mL of streptomycin (Invitrogen). Stable knock-down cell lines were produced by lentiviral transduction of vectors containing the sequence of shRNA targeting the genes of interest and selected with puromycin at a concentration of 10 μg/mL.

### Mycobacterial Strains and Culture

- **Culture Conditions:** The mycobacterial strains used in this study are listed in Extended Table 1. Mycobacteria were grown in Middlebrook 7H9 (Difco) supplemented with 10% OADC (Becton Dickinson), 0.2% glycerol, and 0.05% Tween 80 or tyloxapol (Sigma Aldrich) at 32°C in shaking culture at 150 rpm in the presence of 5 mm glass beads to prevent clumping. Hygromycin was used at a concentration of 100 μg/mL for mCherry/GFP expression, kanamycin was used at a concentration of 50 μg/mL for GFP/DsRed expression, or 25 μg/mL for lux expression. To follow Mm in vitro growth, 2×10^5 bacteria were plated in 200 μl of 7H9 with 10% OADC, 0.05% tyloxapol, and 0.2% glycerol in a 96-well plate. The GFP signal was measured with a Synergy Mx Monochromator-Based Multi-Mode Microplate Reader (Biotek) at 32°C with constant orbital shaking in a black 96 wells plate (PerkinElmer).

### Antibodies, Reagents, Western Blotting, and Immunofluorescence

- **Antibodies and Reagents:** Recombinant nanobodies with the Fc portion of rabbit IgG, which specifically recognize VacA or VacB, were previously characterized^37^. Other antibodies used included pan-vacuolin 221.1.1 (Dr. M. Maniak,^35^), ubiquitin FK2 (Enzo Life Sciences), GFP (pAb from MBL Intl., mAb from Abmart), and Filipin III from *Streptomyces filipinensis* (Sigma-Aldrich). Goat anti-mouse or anti-rabbit IgG coupled to AlexaFluor 488, AlexaFluor 594, AlexaFluor 647 (Invitrogen), or to HRP (Brunschwig) were used as secondary antibodies.
- **Western Blotting:** After SDS-PAGE separation and transfer onto nitrocellulose membranes (Protran), immunodetection was performed as previously described^64^ but with ECL Prime Blocking Reagent (Amersham Biosciences) instead of non-fat dry milk. Detection was performed with ECL Plus (Amersham Biosciences) using a Fusion Fx device (Vilber Lourmat). Quantification of band intensity was performed with ImageJ.
- **Immunofluorescence:** For immunofluorescence, infected Dd cells were fixed with ultra-cold methanol (MeOH) at the indicated time points and immunostained as previously described^65^. BV-2 cells were fixed with 4% PFA in PBS for 20 min and then washed 3 times in PBS before antibody staining. Images were recorded with a Leica SP8 confocal microscope using a 63×1.4 NA oil immersion objectives or Leica Stellaris 8 63×1.4 NA oil immersion objectives. For filipin staining, cells were fixed in a solution of 4% PFA/picric acid as previously described^65^. Then, cells were stained in a solution of 50 μg/ml of filipin in PBS in the dark for 30 min at room temperature. Images were recorded with a spinning disc confocal system (Intelligent Imaging Innovations) mounted on an inverted microscope (Leica DMIRE2; Leica) using the 100×1.4 NA oil objective or Leica Stellaris 8 63×1.4 NA oil immersion objectives.

### Infection Assays

- **General Protocol:** Infections were performed as previously described^8,66^ with few modifications. After infection and phagocytosis, extracellular bacteria were washed off, and attached infected cells were resuspended in filtered HL5c containing a bacteriostatic dose of 5 μg/mL of streptomycin and 5 U/mL of penicillin to prevent growth of extracellular Mm. Mock-infected cells were treated similarly but without bacteria. For CD treatments, the indicated concentration of MβCD (Sigma-Aldrich) or HβCD (Sigma-Aldrich) was added after bacteria phagocytosis to prevent any effect on Mm uptake.
- **BV-2 Infections:** The same protocol was followed for BV-2 infections, with modifications. The day before the infection, 5×10^5 cells/mL were plated in DMEM complemented with FBS. Cells were infected at an MOI of 5 and maintained at 32°C. For CD experiments, prior to infection, the medium was changed to DMEM complemented with delipidated FBS (LRA Sigma #13358-U).

### Phagocytic Plaque Assay

- **Procedure:** Dd plaque formation on a lawn of *Klebsiella pneumoniae* (GE) mixed with mycobacteria was monitored as previously described^67^. A 5×10^8 mycobacteria/mL culture was centrifuged and resuspended in 1.2 mL 7H9 containing a 1:105 dilution of *K. pneumoniae* grown overnight in LB. 50 µL of this suspension were deposited on wells from a 24-well plate containing 2 mL of 7H10-agar (without OADC). Serial dilutions of Dd (10, 10^2, 10^3, or 10^4 cells) were plated onto the bacterial lawns, and plaque formation was monitored after 4-7 days at 25°C.

### Quantitative Real-Time PCR (qRT-PCR)

- **RNA Extraction and cDNA Synthesis:** RNA from mock-infected cells or cells infected with Mm wt, ΔRD1, or *M. smegmatis* was extracted at the indicated time points using the Direct-zol RNA MiniPrep kit (Zymo Research) following the manufacturer’s instructions. 1 μg of RNA was retro transcribed using the iScript cDNA Synthesis Kit and polydT primers (Bio-Rad).
- **PCR Amplification:** The cDNA was amplified using the primers listed in Extended Table 2 and the SsoAdvanced universal SYBR Green supermix (Bio-Rad). Amplimers for vacA, vacB, vacC, and gapdh were detected on a CFX Connect Real-Time PCR Detection System (Bio-Rad). The housekeeping gene gapdh was used for normalization. PCR amplification was followed by a DNA melting curve analysis to confirm the presence of a single amplicon. Relative mRNA levels (2−ΔΔCt) were determined by comparing first the PCR cycle thresholds (Ct) for the gene of interest and gapdh (ΔC) and second Ct values in infected cells vs. mock-infected cells (ΔΔC).

### RNAseq

- **Sample Preparation and Analysis:** Following infection with GFP-expressing bacteria, infected and mock-infected cells were pelleted and resuspended in 500 μl of HL5c, passed through 30 μm filters, and sorted by FACS (Beckman Coulter MoFlo Astrios). The gating was based on cell diameter (forward-scatter) and granularity (side-scatter). Infected (GFP-positive) and non-infected (GFP-negative) sub-fractions were selected based on the GFP intensity (FITC channel). Typically, ∼5 x 10^5 cells of each fraction were collected for RNA isolation. RNA isolation was performed as for qRT-PCR. Quality of RNA libraries, sequencing, and bioinformatic analysis were performed as previously described ^38^.

### Induction of Sterile Damage with LLOMe

- **Experimental Setup:** For LLOMe experiments, 5×10^5 cells were plated in 96-well IBIDI dishes in filtered HL5c. To stain acidic compartments, 1 μM LysoSensor Green DND-189 (ThermoFisher) was added to the cells. After 45 minutes of incubation, excess dye was washed away. Cells were imaged with a spinning disc confocal system (Intelligent Imaging Innovations) mounted on an inverted microscope (Leica DMIRE2; Leica) using the 100×1.4 NA oil objective every minute for 1.5 hours. For quantification of the repair reporters, cell imaging was performed with a 60x water immersion objective with the ImageXpress Micro XL HC microscope every 5 minutes for 1.5 hours using 3 wells per condition with 4 image fields per well with 3 sections per field with a 1 µm interval in three biological replicates of the experiment. Briefly, a first image was taken for all conditions prior to LLOMe addition at the indicated concentration, then the localization of GFP-Vps32 and Alix-GFP was followed. For MβCD treatment, cells were incubated with 2 mM of MβCD 40 minutes prior to LLOMe addition. Analysis was performed per cell and quantified dot formation and area.

### Cytosol-Membrane Separation and rEsxA Incubation

- **Cell Preparation and Fractionation:** 10^9 D. discoideum cells were washed in Sorensen-Sorbitol and resuspended in HESES buffer (20 mM HEPES, 250 mM Sucrose, 5 mM MgCl2, 5 mM ATP) supplemented with protease inhibitors (cOmplete EDTA-free, Roche). Cells were homogenized using a ball homogenizer with a 10 μm clearance. The post-nuclear supernatant was then diluted in HESES buffer and centrifuged at 35,000 rpm in a Sw60 Ti rotor (Beckman) for 1 hour at 4°C. The cytosol (SN, supernatant) and membrane (MB, pellet) fractions were subsequently collected.
- **Protein Incubation and Analysis:** The protein concentration of the cytosol fraction was quantified using the Bradford method. Different quantities of membranes were tested as detailed in Supplementary Figure 6. Ultimately, 400 μg of membranes were incubated with 12 μg of recombinant EsxA (rESAT-6, BEI Resources NR-49424) in HESES Buffer at pH 6 for 20 minutes at room temperature on a wheel. After ultracentrifugation at 45,000 rpm for 1 hour at 4°C in a TLS-55 rotor (Beckman), the membranes were separated into supernatant (SN) and pellet (P) fractions. Equal amounts of SN and P were loaded for western blotting.

### Detergent-Resistant Membrane Isolation

- **Cell Lysis and Fractionation:** 10^8 *D. discoideum* cells were washed in Sorensen-Sorbitol and resuspended in 1 ml of cold Lysis Buffer (50 mM Tris-HCl pH 7.5, 150 mM NaCl, 50 mM Sucrose, 5 mM EDTA, 5 mM ATP, 1 mM DTT) with 1% Triton X-100 supplemented with protease inhibitors. The lysate was incubated at 4°C on a rotating wheel for 30 minutes. After centrifugation at 13,000 rpm for 5 minutes in a tabletop centrifuge at 4°C, the supernatant (Triton Soluble Fraction, TSF) was collected, and the pellet (Triton Insoluble Fraction, TIF) was resuspended in 200 μl of cold Lysis buffer without Triton X-100.
- **Sucrose Gradient Centrifugation:** The TIF was mixed with 800 μl of 80% sucrose (to achieve a final concentration of 65% sucrose) and deposited at the bottom of an ultracentrifuge tube. It was then overlaid with 2 ml of 50% sucrose and 1 ml of 10% sucrose. The sample was centrifuged at 55,000 rpm in a Sw60 Ti rotor (Beckman) for 2 hours at 4°C. The Triton Insoluble Floating Fraction (TIFF) was collected, acetone-precipitated, and resuspended in Laemmli Buffer. Equivalent amounts of each fraction (TSF, TIF, and TIFF) were loaded for western blotting.

---

**Supplementary Figure 1: Endogenous VacA, VacB and sterols are present at the MCV. a.** Representative images of wt cells infected with GFP-expressing Mm wt, fixed at indicated time points and immunostained with recombinants antibodies against endogenous VacA or VacB. Scale bar, 5 μm. **b**. Representative live images of VacC-GFP KI expressing mScarlet-D4H* infected with Mm wt expressing BFP at 6 hpi. Scale bar, 5 μm.

**Supplementary Figure 2: Vacuolin microdomains accumulate at the MCV during infection. a-b**. Indicated non-infected Vac-GFP KI (**a.**) or Vac-OE (**b.**) cells were lysed in cold Triton X-100. The Triton soluble (TSF) and insoluble (TIF) fractions were recovered, as well as the floating fraction (TIFF) from a sucrose gradient. **c-h**. Indicated Vac-GFP KI cells were infected with expressing-mCherry Mm wt and GFP intensity was measured at the indicated time points. **c-h.** Normalized distribution of the indicated Vac-GFP KI intensity during infection and treatment with 2 mM of MβCD at 2 hpi (**c.** and **f.**).and 24 hpi (**d.** and **g.**) (N=3, n≥ 50000 cells). **e.** and **h.** Mean intensity of c-d. and f-g. (mean + sd, N=3, two-way ANOVA, *p≤0.05, **p≤0.01, ***p≤0.005).

**Supplementary Figure 3: Plaque assays with vacuolin KO cells and dose-dependent effect of MβCD. a-b.** Dilutions of wt or vacuolin KO cells were deposited on the lawns of *K. pneumoniae* mixed with indicated mycobacterial. **a.** Representative images of plaque assays taken at day 5. **b.** The plaquing score was determined using a logarithmic scale. (mean ± sem, N=4). **c.** Generation time of GFP-expressing Mm wt monitored for 72 hours in a plate reader (mean fold change ± sd N=3, one-way ANOVA, **p≤0.05). **d.** Wt cells were infected with bioluminescent Mm wt or ΔRD1 and treated with increasing doses of MβCD and luminescence was monitored for 72 hours (mean of the area under the curve ± sd N=3, one-way ANOVA, *p≤0.05, **p≤0.01, ***p≤0.005, ****p≤0.0001). **e.** Wt and ΔvacABC cells were infected with GFP-expressing Mm ΔRD1 and imaged by HC microscopy for 72 hours. The proportion of intracellular and extracellular bacteria was plotted for each time point (mean ± sem, N=3).

**Supplementary Figure 4: Model depicting different possible scenario for increased resistance to infection. a.** Infection course of Mm wt in wt cells. After entry by phagocytosis, Mm hacks phagosome maturation and establishes an MCV. At early stages, Mm induces damage to the MCV membrane, damage is sensed and repaired by the ESCRT and autophagy machineries. When catastrophic level of damage is reached, the escapes to the cytosol where perilipin (Plin) binds their accessible surface. Mm continues to replicate before it egresses and infects neighbouring cells. **b.** Attenuation can be a consequence of no (or less) MCV damage leading to Mm containment in the MCV or phagosome (**b.1.**). Alternatively, attenuation can also result from more efficient damage leading to precocious escape and recapture by the xenophagy machinery (**b.2.**).

**Supplementary Figure 5: Vacuolin KO cells react less and recover faster from LLOMe treatment.** Time-lapse images of movie S1 and S2 of wt and 1′vacABC cells stain with lysosensor and treated with 4.5 mM of LLOMe for a total of 90 min. Insets are magnified 2-fold. Scale bar, 5 μm.

**Supplementary Figure 6: EsxA binds membranes at pH 6, the pH measured in the MCV lumen.** Wt and ΔvacABC cells expressing mCherry-Plin were infected with GFP-expressing Mm wt and treated or not with 2 mM of MβCD, imaged live by HC microscopy at 24 hpi. Plin recruitment around Mm was quantifed (mean + individual replicates, N=4). **b-c.** Recombinant EsxA (rEsxA) was incubated with purified membranes of wt cells at different pH (**b.**) or membrane to peptide ratios (**c.**), before separation in supernatant (SN) and pellet (P) fractions. Identical protein amounts were loaded and immunoblotted with the indicated antibody. **d.** Wt cells were infected with Mm or Mm-L1D labelled with FITC (pH sensitive) and TRITC (pH insensitive). The ratio of the two fluorescence intensities was monitored (mean ± sem, N=2, n≥3). **e.** Recombinant EsxB (rEsxB) was incubated with post-nuclear supernatant (PNS) of wt or ΔvacABC cells, treated or not with 10 mM of MβCD. The PNS was separated into supernatant (SN-cytosol) and pellet (P-membrane) fractions. Identical protein amounts were loaded and immunoblotted with the indicated antibodies.

**Supplementary Figure 7: Disruption of flotillin microdomains confers resistance to infection in mammalian microglial BV-2 cells. a.** Total protein lysates of BV-2 cells, transduced with vectors expressing shRNAs to knock down Flot-1 or Flot2, were subjected to western blot assays using antibodies against Flot-1 or Flot2. As loading control, samples were probed with anti-Importin-β. **b-h.** BV-2 cells were infected with GFP-expressing Mm wt and monitored by HC microscopy for 48 hours. **b.** Representative images of bacteria at high or low brightness and contrast. **c**. Scheme illustrating the microcolony morphology. **d-e**. Violin plots of the distribution of microcolony area at 3 hpi (**d.**) and 33 hpi (**e.**) (Median, and quantiles; one representative experiment). Dashed line represents the median of the control condition, BV-2 wt cells infected with Mm wt. **f-h.** Median of microcolony area of **d-e** are plotted as a time course for each condition and surrounded by the lower and higher quantiles.

**Supplementary Movie S1: Time-lapse movie of wt cells stained with lysosensor and treated with LLOMe**

**Supplementary Movie S2: Time-lapse movie of 1′vacABC cells stained with lysosensor and treated with LLOMe.**

## Notes

### Competing Interest Statement

The authors have declared no competing interest.

## REFERENCES

1. World Health Organization. Global tuberculosis report 2023. (2023).

2. Russell, D. G. Who puts the tubercle in tuberculosis? Nat. Rev. Microbiol. 5, 39–47 (2007).

3. Russell, D. G. Mycobacterium tuberculosis: here today, and here tomorrow. Nat. Rev. Mol. Cell Biol. 2, 569–586 (2001).

4. Stinear, T. P. et al. Insights from the complete genome sequence of Mycobacterium marinum on the evolution of Mycobacterium tuberculosis. Genome Res. 18, 729–741 (2008).

5. Tobin, D. M. & Ramakrishnan, L. Comparative pathogenesis of Mycobacterium marinum and Mycobacterium tuberculosis. Cell. Microbiol. 10, 1027–1039 (2008).

6. Boulais, J. et al. Molecular characterization of the evolution of phagosomes. Mol. Syst. Biol. 6, 423 (2010).

7. Solomon, J. M., Leung, G. S. & Isberg, R. R. Intracellular Replication of Mycobacterium marinum within Dictyostelium discoideum: Efficient Replication in the Absence of Host Coronin. Infect. Immun. 71, 3578–3586 (2003).

8. Hagedorn, M. & Soldati, T. Flotillin and RacH modulate the intracellular immunity of Dictyostelium to Mycobacterium marinum infection. Cell. Microbiol. 9, 2716–2733 (2007).

9. Cosson, P. & Soldati, T. Eat, kill or die: when amoeba meets bacteria. Curr. Opin. Microbiol. 11, 271–276 (2008).

10. Eichinger, L. et al. The genome of the social amoeba Dictyostelium discoideum. Nature 435, 43– 57 (2005).

11. Dunn, J. D. et al. Eat Prey, Live: Dictyostelium discoideum As a Model for Cell-Autonomous Defenses. Front. Immunol. 8, 1906 (2018).

12. Guallar-Garrido, S. & Soldati, T. Exploring host–pathogen interactions in the Dictyostelium discoideum–Mycobacterium marinum infection model of tuberculosis. (2024).

13. Cosson, P. & Lima, W. C. Intracellular killing of bacteria: is Dictyostelium a model macrophage or an alien? Cell. Microbiol. 16, 816–823 (2014).

14. Barisch, C., Kalinina, V., Lefrancois, L. H., Appiah, J. & Soldati, T. Think zinc: Role of zinc poisoning in the intraphagosomal killing of bacteria by the amoeba Dictyostelium. bioRxiv356949 (2018) doi:10.1101/356949.

15. Hanna, N. et al. Zn ^2+^ Intoxication of Mycobacterium marinum during Dictyostelium discoideum Infection Is Counteracted by Induction of the Pathogen Zn ^2+^ Exporter CtpC. mBio 12, e01313–20 (2021).

16. Lewis, K. N. et al. Deletion of RD1 from Mycobacterium tuberculosis Mimics Bacille Calmette-Guérin Attenuation. J. Infect. Dis. 187, 117–123 (2003).

17. Gao, L. et al. A mycobacterial virulence gene cluster extending RD1 is required for cytolysis, bacterial spreading and ESAT-6 secretion. Mol. Microbiol. 53, 1677–1693 (2004).

18. Cardenal-Muñoz, E., Barisch, C., Lefrançois, L. H., López-Jiménez, A. T. & Soldati, T. When Dicty Met Myco, a (Not So) Romantic Story about One Amoeba and Its Intracellular Pathogen. Front. Cell. Infect. Microbiol. 7, 529 (2018).

19. Cardenal-Muñoz, E. et al. Mycobacterium marinum antagonistically induces an autophagic response while repressing the autophagic flux in a TORC1- and ESX-1-dependent manner. PLOS Pathog. 13, e1006344 (2017).

20. López-Jiménez, A. T. et al. The ESCRT and autophagy machineries cooperate to repair ESX-1-dependent damage at the Mycobacterium-containing vacuole but have opposite impact on containing the infection. PLOS Pathog. 14, e1007501 (2018).

21. Raykov, L., Mottet, M., Nitschke, J. & Soldati, T. A TRAF-like E3 ubiquitin ligase TrafE coordinates ESCRT and autophagy in endolysosomal damage response and cell-autonomous immunity to Mycobacterium marinum. eLife 12, e85727 (2023).

22. Bernard, E. M., et al. *M. tuberculosis* infection of human iPSDM reveals complex membrane dynamics during xenophagy evasion. J. Cell Sci. jcs.252973 (2020) doi:10.1242/jcs.252973.

23. Augenstreich, J. et al. ESX-1 and phthiocerol dimycocerosates of *Mycobacterium tuberculosis* act in concert to cause phagosomal rupture and host cell apoptosis. Cell. Microbiol. 19, e12726 (2017).

24. Cambier, C., Banik, S., Buonomo, J. A. & Bertozzi, C. Spreading of a Virulence Lipid into Host Membranes Promotes Mycobacterial Pathogenesis. http://biorxiv.org/lookup/doi/10.1101/845081 (2019) doi:10.1101/845081.

25. Lerner, T. R. et al. Phthiocerol dimycocerosates promote access to the cytosol and intracellular burden of Mycobacterium tuberculosis in lymphatic endothelial cells. BMC Biol. 16, 1 (2018).

26. Dermine, J.-F. et al. Flotillin-1-enriched Lipid Raft Domains Accumulate on Maturing Phagosomes. J. Biol. Chem. 276, 18507–18512 (2001).

27. Morrow, I. C. & Parton, R. G. Flotillins and the PHB Domain Protein Family: Rafts, Worms and Anaesthetics: Flotillins and the PHB Family. Traffic 6, 725–740 (2005).

28. Otto, G. P. & Nichols, B. J. The roles of flotillin microdomains - endocytosis and beyond. J. Cell Sci. 124, 3933–3940 (2011).

29. Corkery, D. P., et al. *Vibrio cholerae* cytotoxin MakA induces noncanonical autophagy resulting in the spatial inhibition of canonical autophagy. J. Cell Sci. jcs.252015 (2020) doi:10.1242/jcs.252015.

30. Jia, X., et al. *V. cholerae* MakA is a cholesterol-binding pore-forming toxin that induces non-canonical autophagy. J. Cell Biol. 221, e202206040 (2022).

31. Portnoy, D. A., Auerbuch, V. & Glomski, I. J. The cell biology of Listeria monocytogenes infection. J. Cell Biol. 158, 409–414 (2002).

32. Arellano-Reynoso, B. et al. Cyclic β-1,2-glucan is a brucella virulence factor required for intracellular survival. Nat. Immunol. 6, 618–625 (2005).

33. Korhonen, J. T. et al. Flotillin-1 (Reggie-2) Contributes to Chlamydia pneumoniae Growth and Is Associated with Bacterial Inclusion. Infect. Immun. 80, 1072–1078 (2012).

34. Xiong, Q., Lin, M., Huang, W. & Rikihisa, Y. Infection by Anaplasma phagocytophilum Requires Recruitment of Low-Density Lipoprotein Cholesterol by Flotillins. mBio 10, e02783–18 (2019).

35. Jenne, N., Rauchenberger, R., Hacker, U., Kast, T. & Maniak, M. Targeted gene disruption reveals a role for vacuolin B in the late endocytic pathway and exocytosis. J. Cell Sci. 111, 61–70 (1998).

36. Wienke, D., Drengk, A., Schmauch, C., Jenne, N. & Maniak, M. Vacuolin, a flotillin/reggie-related protein from Dictyostelium oligomerizes for endosome association. Eur. J. Cell Biol. 85, 991–1000 (2006).

37. Bosmani, C. et al. Vacuolins and myosin VII are required for phagocytic uptake and phagosomal membrane recycling in Dictyostelium discoideum. J. Cell Sci. 133, (2020).

38. Hanna, N., et al. *Time-Resolved RNA-Seq Profiling of the Infection of* Dictyostelium Discoideum *by* Mycobacterium Marinum *Reveals an Integrated Host Response to Damage and Stress*. http://biorxiv.org/lookup/doi/10.1101/590810 (2019) doi:10.1101/590810.

39. Stajdohar, M. et al. dictyExpress: a web-based platform for sequence data management and analytics in Dictyostelium and beyond. BMC Bioinformatics 18, 291 (2017).

40. Lamrabet, O., Jauslin, T., Lima, W. C., Leippe, M. & Cosson, P. The multifarious lysozyme arsenal of Dictyostelium discoideum. Dev. Comp. Immunol. 107, 103645 (2020).

41. Marek, M., Vincenzetti, V. & Martin, S. G. Sterol biosensor reveals LAM-family Ltc1-dependent sterol flow to endosomes upon Arp2/3 inhibition. J. Cell Biol. 219, e202001147 (2020).

42. Janssen, K.-P., Rost, R., Eichinger, L. & Schleicher, M. Characterization of CD36/LIMPII Homologues inDictyostelium discoideum. J. Biol. Chem. 276, 38899–38910 (2001).

43. Clark, H. F. & Shepard, C. C. EFFECT OF ENVIRONMENTAL TEMPERATURES ON INFECTION WITH *MYCOBACTERIUM MARINUM* (BALNEI) OF MICE AND A NUMBER OF POIKILOTHERMIC SPECIES. J. Bacteriol. 86, 1057–1069 (1963).

44. Fey, P., Kowal, A. S., Gaudet, P., Pilcher, K. E. & Chisholm, R. L. Protocols for growth and development of Dictyostelium discoideum. Nat. Protoc. 2, 1307–1316 (2007).

45. Hagedorn, M., Rohde, K. H., Russell, D. G. & Soldati, T. Infection by tubercular mycobacteria is spread by nonlytic ejection from their amoeba hosts. Science 323, 1729–1733 (2009).

46. Mittal, E. et al. Mycobacterium tuberculosis Type VII Secretion System Effectors Differentially Impact the ESCRT Endomembrane Damage Response. mBio 9, e01765–18 (2018).

47. Bussi, C. et al. Stress granules plug and stabilize damaged endolysosomal membranes. Nature 623, 1062–1069 (2023).

48. Vietri, M., Radulovic, M. & Stenmark, H. The many functions of ESCRTs. Nat. Rev. Mol. Cell Biol. 21, 25–42 (2020).

49. Barisch, C., Paschke, P., Hagedorn, M., Maniak, M. & Soldati, T. Lipid droplet dynamics at early stages of *M ycobacterium marinum* infection in *D ictyostelium*: Lipid droplets in mycobacterium infection. Cell. Microbiol. 17, 1332–1349 (2015).

50. de Jonge, M. I. et al. ESAT-6 from Mycobacterium tuberculosis Dissociates from Its Putative Chaperone CFP-10 under Acidic Conditions and Exhibits Membrane-Lysing Activity. J. Bacteriol. 189, 6028–6034 (2007).

51. Koiri, D. et al. Real-time visualization reveals Mycobacterium tuberculosis ESAT-6 disrupts phagosome via fibril-mediated vesiculation. Preprint at 10.1101/2024.04.19.590309 (2024).

52. Ray, S., Vazquez Reyes, S., Xiao, C. & Sun, J. Effects of membrane lipid composition on Mycobacterium tuberculosis EsxA membrane insertion: A dual play of fluidity and charge. Tuberculosis 118, 101854 (2019).

53. Golovkine, G. R. et al. Autophagy restricts Mycobacterium tuberculosis during acute infection in mice. Nat. Microbiol. 8, 819–832 (2023).

54. Wang, F. et al. ATG5 provides host protection acting as a switch in the atg8ylation cascade between autophagy and secretion. Dev. Cell 58, 866–884.e8 (2023).

55. Guého, A., Bosmani, C., Nitschke, J. & Soldati, T. Proteomic characterization of the *Mycobacterium marinum* -containing vacuole in *Dictyostelium discoideum*. Preprint at 10.1101/592717 (2019).

56. Machelart, A. et al. Intrinsic Antibacterial Activity of Nanoparticles Made of β-Cyclodextrins Potentiates Their Effect as Drug Nanocarriers against Tuberculosis. ACS Nano 13, 3992–4007 (2019).

57. Astarie-Dequeker, C. et al. Phthiocerol Dimycocerosates of M. tuberculosis Participate in Macrophage Invasion by Inducing Changes in the Organization of Plasma Membrane Lipids. PLOS Pathog. 5, e1000289 (2009).

58. Leon, J. D. et al. Mycobacterium tuberculosis ESAT-6 exhibits a unique membrane-interacting activity that is not found in its ortholog from non-pathogenic Mycobacterium smegmatis. J. Biol. Chem. jbc.M112.420869 (2012) doi:10.1074/jbc.M112.420869.

59. Augenstreich, J. & Briken, V. Host Cell Targets of Released Lipid and Secreted Protein Effectors of Mycobacterium tuberculosis. Front. Cell. Infect. Microbiol. 10, 595029 (2020).

60. Aguilera, J. et al. N α-Acetylation of the virulence factor EsxA is required for mycobacterial cytosolic translocation and virulence. J. Biol. Chem. 295, 5785–5794 (2020).

61. Cambier, C., Banik, S. M., Buonomo, J. A. & Bertozzi, C. R. Spreading of a mycobacterial cell-surface lipid into host epithelial membranes promotes infectivity. eLife 9, e60648 (2020).

62. Barisch, C. & Soldati, T. Breaking fat! How mycobacteria and other intracellular pathogens manipulate host lipid droplets. Biochimie (2017) doi:10.1016/j.biochi.2017.06.001.

63. Foulon, M., Listian, S. A., Soldati, T. & Barisch, C. Conserved mechanisms drive host-lipid access, import, and utilization in Mycobacterium tuberculosis and M. marinum. in Biology of Mycobacterial Lipids 133–161 (Elsevier, 2022). doi:10.1016/B978-0-323-91948-7.00011-7.

64. Schwarz, E. C., Neuhaus, E. M., Kistler, C., Henkel, A. W. & Soldati, T. Dictyostelium myosin IK is involved in the maintenance of cortical tension and affects motility and phagocytosis. J Cell Sci 113, 621–633 (2000).

65. Hagedorn, M., Neuhaus, E. M. & Soldati, T. Optimized Fixation and Immunofluorescence Staining Methods for *Dictyostelium* Cells. in Dictyostelium discoideum Protocols vol. 346 327–338 (Humana Press, New Jersey, 2006).

66. Arafah, S. et al. Setting up and monitoring an infection of Dictyostelium discoideum with mycobacteria. Methods Mol. Biol. Clifton NJ 983, 403–417 (2013).

67. Alibaud, L. et al. A Mycobacterium marinum TesA mutant defective for major cell wall-associated lipids is highly attenuated in Dictyostelium discoideum and zebrafish embryos. Mol. Microbiol. 80, 919–934 (2011).

