## Supplementary Figures for "Membrane microdomains are crucial for *Mycobacterium marinum* EsxA-dependent membrane damage, escape to the cytosol and infection"

Bosmani and Perret et al.,
Supplementary figure 1

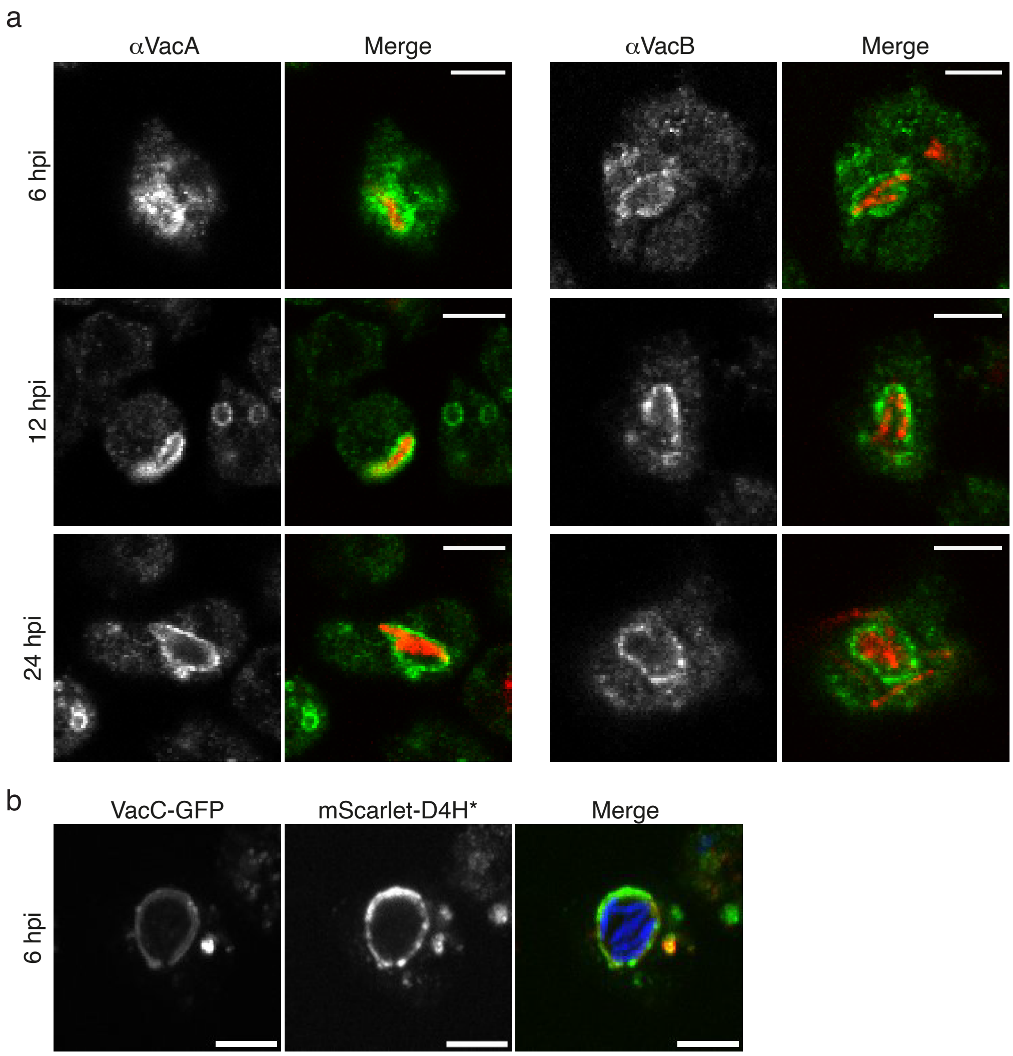

Bosmani and Perret et al.,
Supplementary figure 2

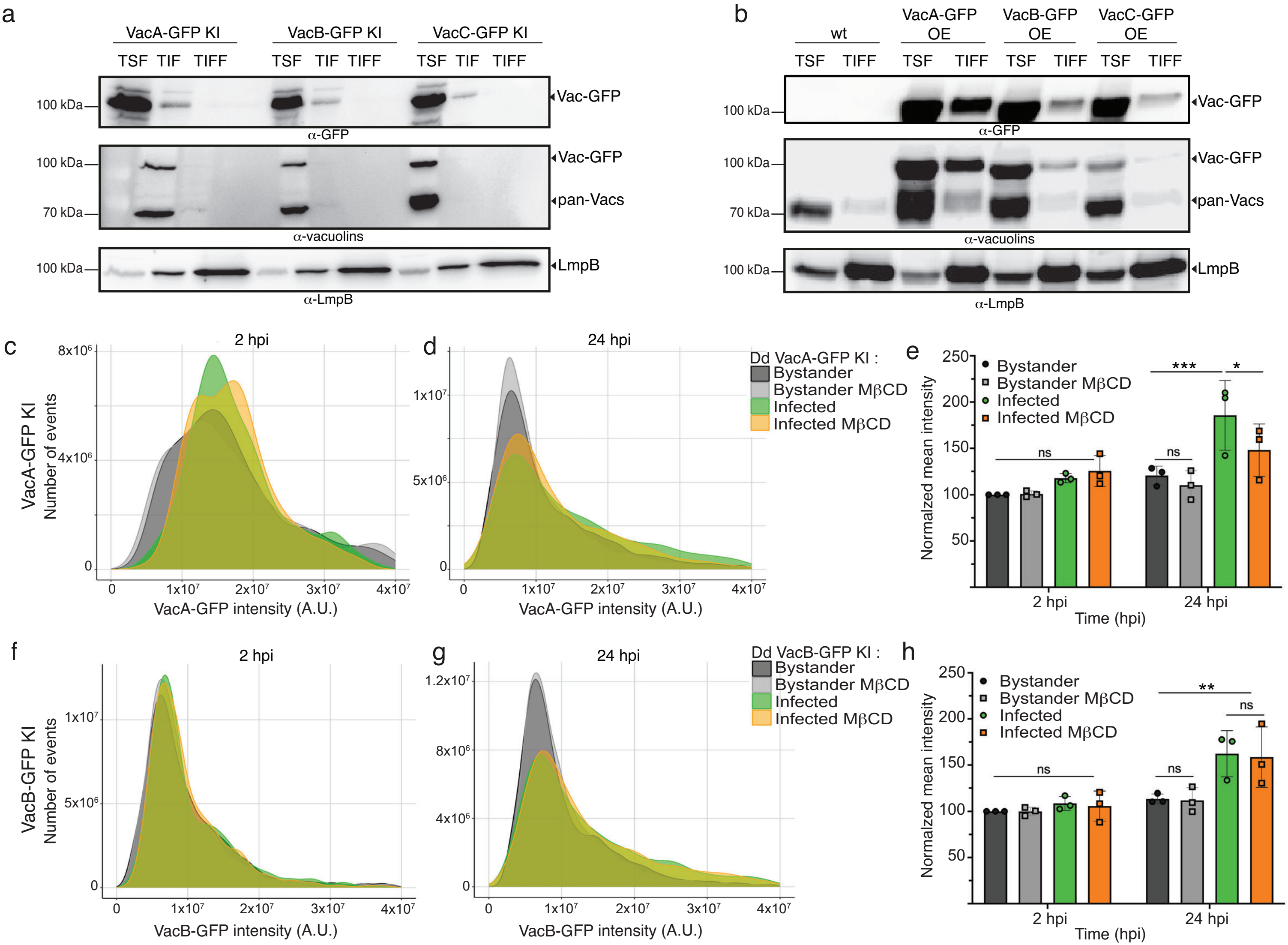

Bosmani and Perret et al.,
Supplementary figure 3

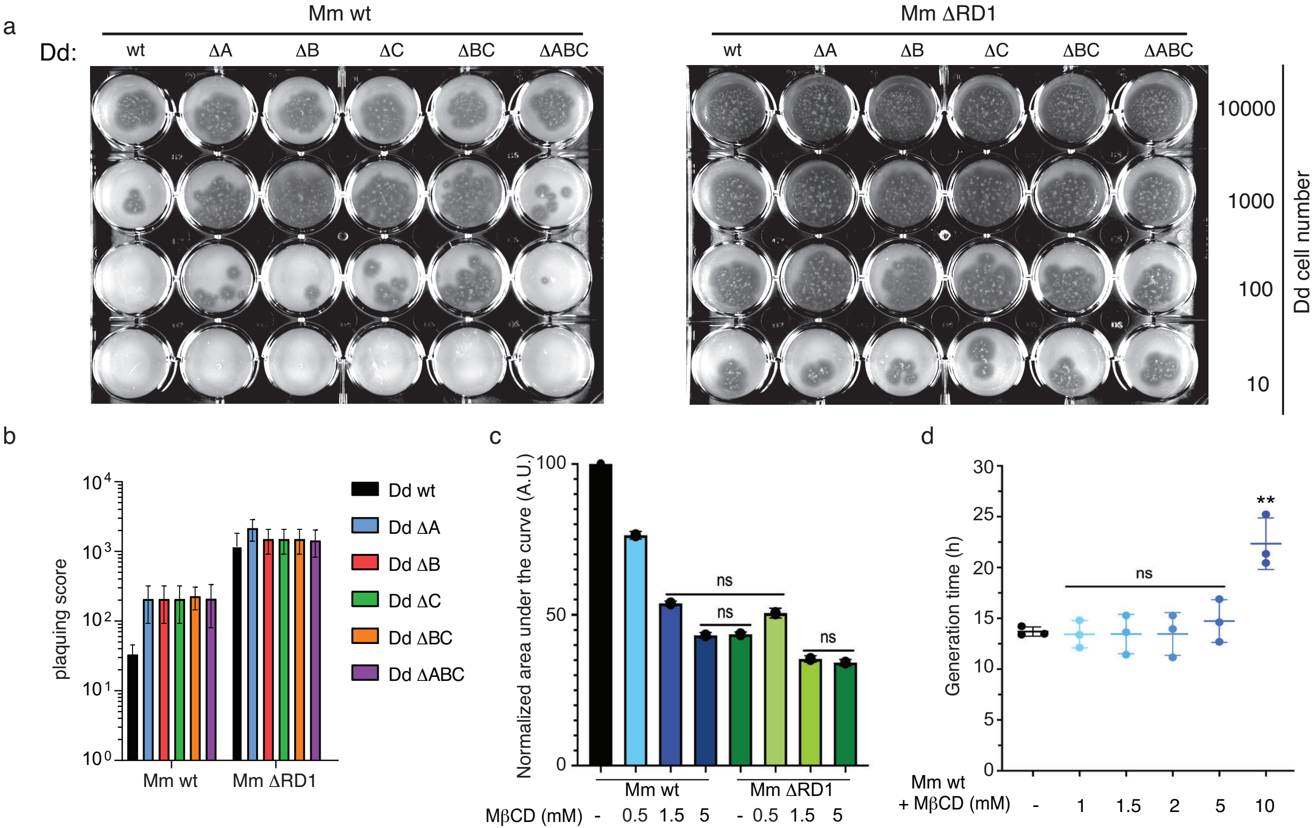

Bosmani and Perret et al.,
Supplementary figure 4

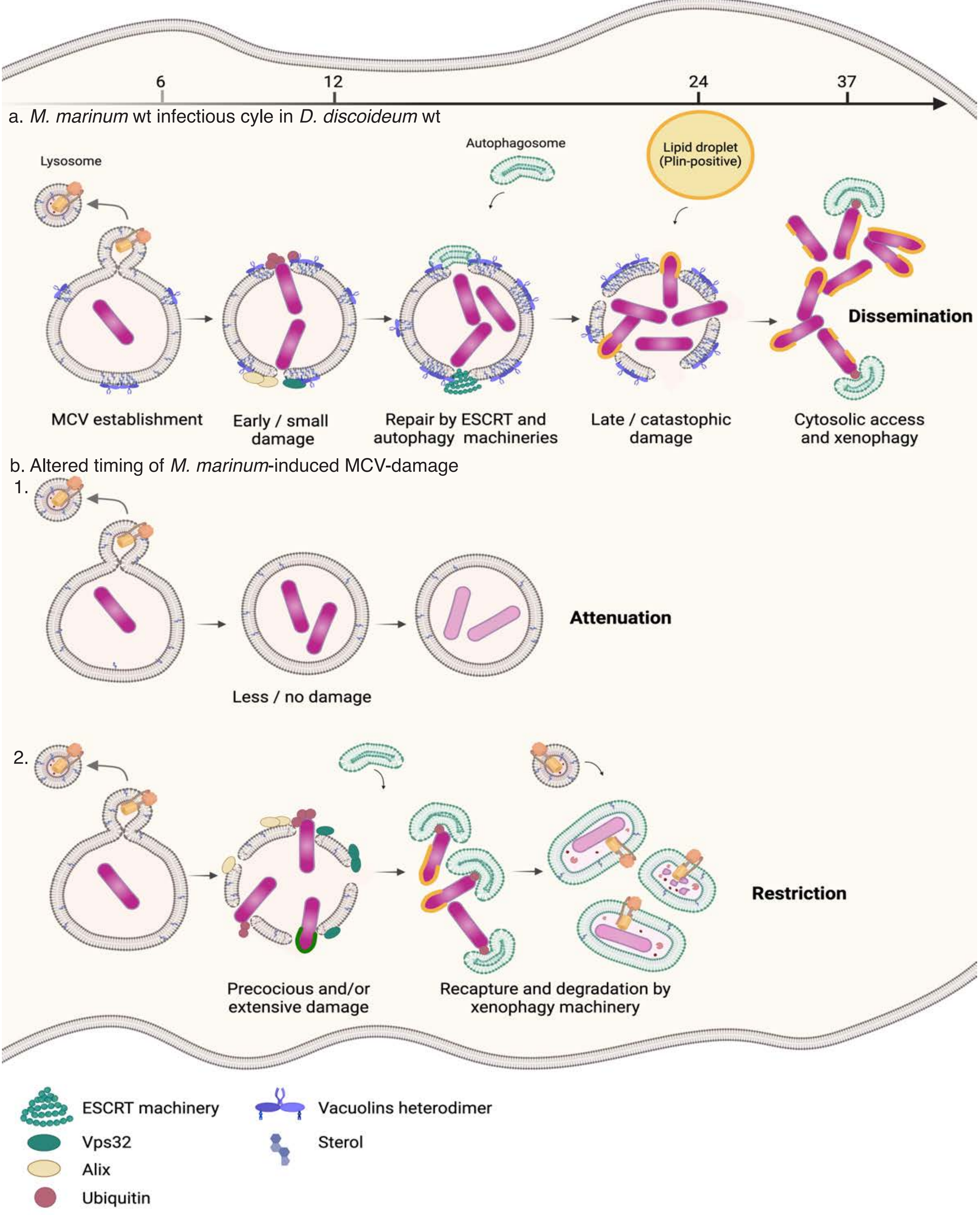

Bosmani and Perret et al.,
Supplementary figure 5

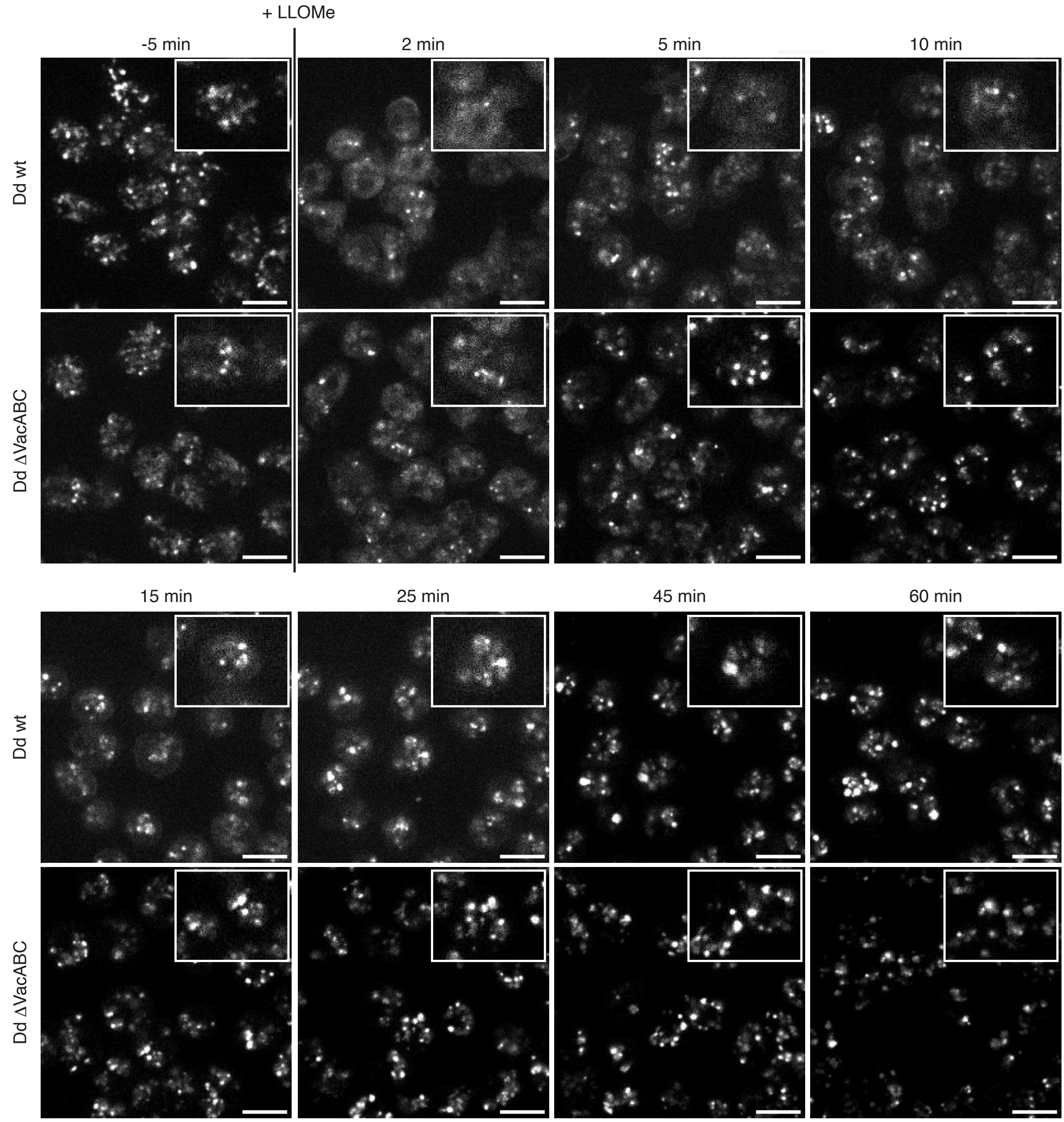

Bosmani and Perret et al.,
Supplementary figure 6

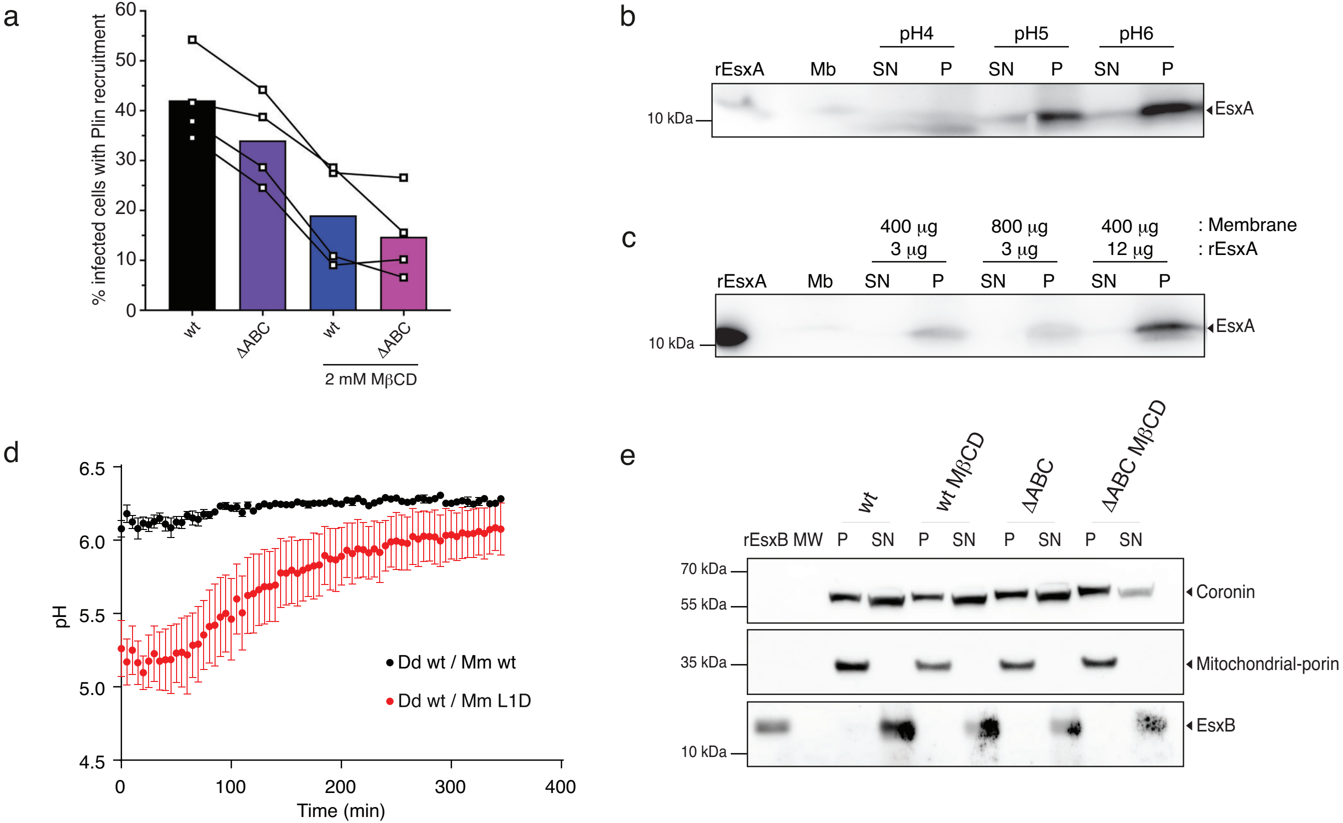

Bosmani and Perret et al.,
Supplementary figure 7

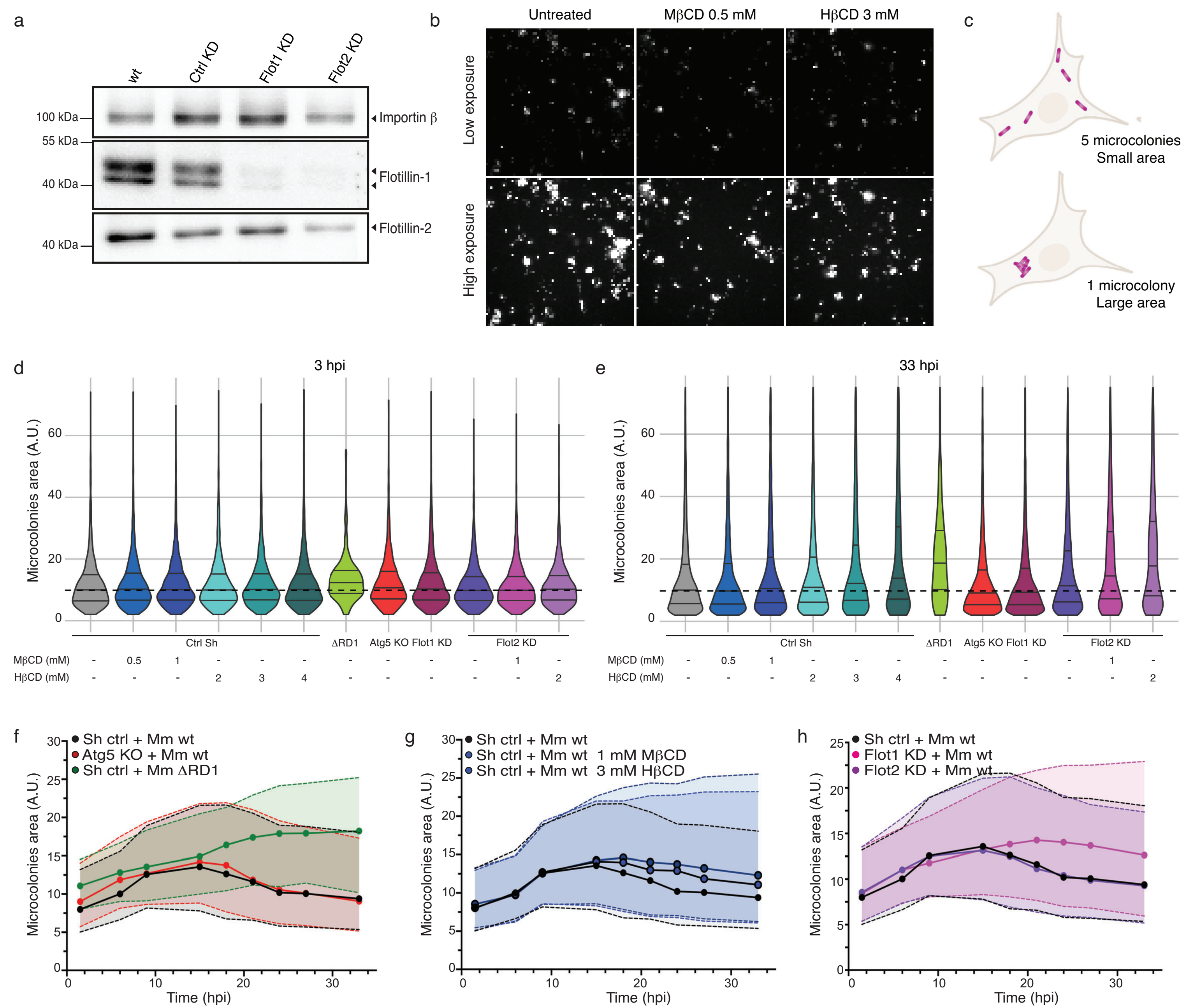
